# UhAVR1, an HR-triggering avirulence effector of *Ustilago hordei*, is secreted via the ER-Golgi pathway to the cytosol of barley coleoptile cells and contributes to virulence early in infection

**DOI:** 10.1101/2020.08.17.254789

**Authors:** Ana Priscilla Montenegro Alonso, Shawkat Ali, Xiao Song, Rob Linning, Guus Bakkeren

**Affiliations:** Department of Botany, University of British Columbia, Vancouver, BC V6T 1Z4, Canada; Agriculture and Agri-Food Canada, Kentville Research and Development Centre, Kentville, NS B4N 1J5, Canada; Sandstone Pharmacies Glenmore Landing Calgary-Compounding, 167D, 1600 – 90 Ave SW Calgary, AB T2V 5A8, Canada; Agriculture and Agri-Food Canada, Summerland Research and Development Centre, Summerland, BC V0H 1Z0, Canada

**Keywords:** gene-for-gene interaction, resistance gene, *Hordeum vulgare*, hypersensitive response, foxtail mosaic virus, FoMV, brefeldin A

## Abstract

The basidiomycete *Ustilago hordei* (*Uh*) causes covered smut disease of barley and oats. Virulence effectors that aid the infection process and support the pathogen’s lifestyle have been described for this fungus. Genetically, six avirulence genes are known and one codes for UhAVR1, the only proven avirulence effector identified in smut pathogens to date that triggers complete immunity in barley cultivars carrying the resistance gene *Ruh1*. A prerequisite for resistance breeding is understanding the host targets and molecular function of UhAVR1. Analysis of this effector upon natural infection of barley coleoptiles using teliospores showed that UhAVR1 is expressed during the early stages of fungal infection where it leads to HR triggering in resistant cultivars or performs its virulence function in susceptible cultivars. Fungal secretion of UhAVR1 is directed by its signal peptide and occurs via the BrefeldinA-sensitive ER-Golgi pathway, both in cell culture away from its host, and during barley interaction. Transient expression of this effector in barley and a heterologous host, *Nicotiana benthamiana* (*Nb*), supports a cytosolic localization. Delivery of UhAVR1 via foxtail mosaic virus, *Pseudomonas* species or *Agrobacterium-mediated* suppression of cell inducers in barley and *Nb* support a role in the suppression of a common component(s) of ETI and PTI which is conserved in both plant systems.

## 1. Introduction

The smut fungus *Ustilago hordei* (*Uh*) is a facultative biotroph responsible for causing covered smut disease of barley and oats. The infection process of this fungus starts when teliospores contaminating the seed hull of barley, germinate together with the seed, produce a promycelium which, through meiosis, produce haploid basidiospores of two different mating types, *MAT-1* and *MAT-2*. Basidiospores of opposite mating type induce in each other conjugation hyphae which merge to initiate mating [1]. This process results in the production of dikaryotic hyphae which penetrate the epidermal cells of barley seedlings [2–4]. Intra- and inter-cellular growth in the host allows the fungus to establish in the meristematic tissue where it will persist until floral differentiation [3,5]. There, after nuclear fusion, diploid teliospores will mature and replace the seed kernels or form pustules on the flag leaf [3,6]. The infection cycle is determined by the developmental stages of the host plant and takes 2 to 3 months without the appearance of visible symptoms until the emergence of flag leaves and smutted seed heads [7,8].

Plant colonizing microbes, independent of their lifestyles, secrete effector proteins to influence the interaction with their host [9]. These effectors have different modes of actions and expression patterns to support the pathogens’ lifestyle, either by modulating the plant immune system or by manipulating the host physiology to support the microbes needs, leading to compatibility [9–11]. To counteract the action of pathogens, plants possess surveillance systems that recognize and neutralize the invading microbes leading to incompatible interactions [9,10].

Effectors or their activities which are recognized by a corresponding resistance gene in a host are known as avirulence effectors [12]. Genome analysis of *Uh* revealed 333 predicted candidate secreted effectors while six avirulence genes have been described genetically [8,13–15]. *UhAvr1* (UHOR_10022, NCBI protein ID CCF49778.1) is the only proven avirulence effector that has been identified in smut pathogens [16]. A gene family formed by *UhAvr1* and its paralogous gene (UHOR_10021) encompasses five genes in *Ustilago maydis* (*Um*), three in *Sporisorium reilianum*, six in *Sporisorium scitamineum* and only one in the related epiphytic fungus *Moesziomyces albugensi* [16,17]. Recently, four effectors (*Uvi1-Uvi4*) contributing to virulence in *Uh* have been identified by gene deletion assays [18].

Incompatibility in the *Uh*-barley pathosystem follows a typical ‘gene for gene’ interaction where the presence of a genetic avirulence factor in a specific *Uh* race and a matching resistance gene in a barley cultivar leads to immunity [19]. Fungal races possessing avirulence gene *UhAvr1* trigger a microscopic HR response after penetration and interaction with the host cytoplasm of cultivars harbouring the corresponding resistance gene *Ruh1* [2]. To date, six resistance genes have been identified in barley, however only resistance gene *Ruh1* has been genetically mapped; this gene is found on barley chromosome 7H [14,15,19,20].

The goal of this study was to investigate how UhAVR1 is being secreted, its localization in the host cell and its potential role during compatible and incompatible interactions. Our results indicate that the effector UhAVR1 is being secreted via the conventional ER-Golgi secretory pathway of *Uh*, directed by the action of its signal peptide, and is found in the cytosol of barley cells. *UhAvr1* is expressed during the first days of infection of barley suggesting its importance during the initial fungal establishment in the plant when it bestows a virulence effect on the fungus. In addition, expression of UhAVR1 using foxtail mosaic virus (FoMV), *Pseudomonas* bacteria and co-expression of this effector with cell death inducers in *Nicotiana benthamiana* and barley plants, suggests that this effector suppresses component/s of plant immunity that are conserved in both plants systems.

## 2. Materials and Methods

### 2.1. Biological materials and growth conditions

Two barley cultivars, cv. Odessa (*ruh1*, universal susceptible) and cv. Hannchen (*Ruh1*), were used in this study. Barley seeds were dehulled and surface sterilized as described previously [16]. Both barley cultivars were grown at 23°C and 16 hours light in environmentally controlled growth chambers (Conviron) unless stated otherwise. For FoMV assays, barley was grown following conditions as described [21]. For this assay, an additional barley line SM89010 (*Ruh1*) was also used [20]. *Nicotiana benthamiana* (*Nb*) plants were grown under similar conditions as the barley.

*Uh* strains were cultured at 22°C in complete medium (CM) [22] with appropriate antibiotic when needed: 5 μg/ml of Carboxin (Sigma-Aldrich) or 40 μg/ml of Zeocin (Invitrogen). All fungal strains used in this work are described in Table S1. *Pseudomonas syringae* pv. *atropurpurea* (*Psa*) isolate 1304, a pathogen of Italian ryegrass [23], was cultured in Luria-Bertani (LB) medium at 28°C with the appropriate antibiotics.

### 2.2. Generation of Ustilago hordei strains

To generate Uh1398 (*MAT-1 ΔUhAvr1* [*SP:GFP:UhAvr1*]), fragment SP:GFP:*UhAvr1:*Nos was synthesized (Bio Basic Inc., ON, Canada) and cloned via restriction digestion into a previously described backbone vector named as replacement construct [16] (Figure S1A). The final construct was transformed into *E. coli* and verified by sequencing. The construct was then linearized by SpeI and used for homologous transformation into strain Uh1289 (*MAT-1 ΔUhAvr1*) placing *SP:GFP:UhAvr1* in the *UhAvr1* endogenous location. Positive transformants were confirmed by performing DNA blot analysis using 5 μg of genomic DNA as in [16]. A 1.3 kb probe that binds to the 3’ flank region of *UhAvr1* was amplified from Uh364 gDNA using primers 1794 + 1795 (all primer sequences are given in Table S2), digested with SalI and BglII, followed by labelling with *α*-P^32^ dCTP using the random primer labelling system (Life Technology). A total of three strains, originated from three independent events, expressing SP:GFP:UhAvr1 (Uh1397, Uh1398 and Uh1399) were obtained and used in subsequent experiments.

### 2.3. Generation of the FunGus secretion system

The new fungal secretion system (FunGus) utilizes an episomal plasmid that can be expressed in *Uh* cells. This episomal plasmid contains a GateWay™ (Invitrogen) recombineering cassette that allows the cloning and expression of fungal effectors without a signal peptide (SP). The *UhAvr1* SP present on the episomal plasmid backbone allows the delivery of effectors. An interim vector was prepared by synthesizing a 1491-bp DNA fragment containing the *UhAvr1* promoter:SP:EcoRV:3HA:STOP:UhAvr1 terminator and flanked by BamHI restriction sites (Life Technologies Inc., Burlington, ON, Canada). The fragment was cloned into the unique BamHI site of *Ustilago*-specific vector pCM60 [24] which has an *Um*-specific Hsp70 promoter-driven hygromycin B resistance cassette. For constitutive high expression, we replaced the *UhAvr1* promoter with the UmHsp70 promoter while leaving the *UhAvr1* SP. To this end, an 836 bp synthetic gene fragment Spe1-UmHsp70 promoter:UhAvr1 SP-EcoRV replaced the Spe1-UhAvr1 promoter:UhAvr1 SP-EcoRV part. Since two UmHsp70 promoter sequences in the same construct may cause instability, the UmHsp70 hygromycin B resistance cassette needed to be replaced. Therefore, pCM60 was modified to delete the hygromycin cassette through Nar1 digestion, blunt end formation by the Klenow fragment of DNA polymerase I, a second digestion with Sph1 and blunt end formation by T4 polymerase, and self-ligation. This interim vector was subsequently digested with PvuII and Xma1, after which a 1.9 kb Xma1-EcoRV fragment from plasmid pDONR-CbxR, harbouring the carboxin resistance gene [25], was inserted to give plasmid pCM100 a unique BamHI site. The complete BamH1-HSP70 promoter:UhAvr1 SP-EcoRV-3HA tag:STOP:UhAvr1 terminator-BamHI fragment was inserted and the GateWay™ reading frame A cassette (Thermo Fisher Scientific) added in the unique EcoRV site. This allows easy in-frame integration via recombineering of any effector cloned in pENTR/D-TOPO. By not including a stop codon in the inserting effector, a triple hemagglutinin (3xHA) tag would be added to allow for epitope tagging and protein expression studies. This gave construct pUHESdest (plasmid Ustilago High Effector Secretion destination vector; Figure 5A).

An entry vector containing UhAvr1-SP:mCherry+STOP was created by PCR amplification with primers 1248 + 2034 followed by GateWay™ TOPO cloning. The GateWay™ LR reaction of this newly generated entry vector was performed with pUHESdest, resulting in pUHES:UhAvr1+SP:mCherry (Figure S1B). A SP-deletion mutant, pUHES:ΔSP:UhAvr1-SP:mCherry was generated by inverse PCR amplification of pUHES:UhAvr1+SP:mCherry using primers 2098 + 2099 which removed its signal peptide. These primers contained unique EcoRI sites that allowed the circularization of the vector (Figure S1B). pUHES:UmPit2+SP:mCherry with its natural signal peptide was generated via NEBuilder HiFi DNA Assembly Cloning Kit (NEB) by combining PCR-amplified UmPit2+SP:mCherry using primers 2127 + 2128 and PCR-amplified pUHESdest using primers 2125 + 2126 without *UhAvr1* signal peptide, GateWay™ cassette and 3xHA tag (Figure S1B). All generated constructs were verified by sequencing and transformed into Uh1351 (*MAT-1 UhAvr1 [otef:gfp]*) as in [16]. The generated strains (Uh1430, Uh1434 and Uh1440) were further used for *in vitro* mating experiments as in [3], for mating tests as in [1], for confocal microscopy, and protein blots.

### 2.4. Transient expression assays

An entry vector (pENTR/D-SPUhAvr1-STOP) with the *UhAvr1* SP was generated by PCR amplification from *UhAvr1* harbouring entry clones using primers 1247 + 2035 followed by the GateWay™ TOPO cloning reaction. The construct consisted of 19 amino acids of UhAVR1 SP predicted by SignalP [26] plus an additional six amino acids downstream of the predicted signal peptide cleavage site. The GateWay™ LR reaction was performed between this generated entry clone and vector pK7FWG2 resulting in 35S:SP:GFP (Figure S1C). Similarly, 35S:UhAvr1-SP:GFP and 35S:UhAvr1+SP:GFP (Figure S1C) were generated by GateWay™ LR reactions between the UhAvr1-SP or UhAvr1+SP entry vectors and the pK7FWG2 destination vector [27].

To generate 35S:UhAvr1+SP:mCherry (Figure S1D), UhAvr1+SP was amplified by PCR using primers 2031 + 2032 with added BamHI sites that allowed restriction cloning into pCAMBIA:mCherry. Whereas, to prepare 35S:UmPit2+SP:mCherry (Figure S1D), UmPit2+SP (UMAG_01375, NCBI protein ID XP_011387264.1, Table S3) was amplified from cDNA of 10 day-old Golden Bantam maize seedlings infected with mixed cultures of *Um* (Um001 and Um002) as in [28]. The PCR primers 2129 + 2130 which contained BamHI sites, allowed ligation of the amplicon into the single BamHI site of pCAMBIA:mCherry. As positive fluorescent controls, 35S:GFP in binary vector pBIN+ [29] and 35S:mCherry in vector pCAMBIA [30] (Figure S1D) were used. All constructs were sequenced and transformed into *A. tumefaciens* GV3101. Furthermore, constructs used for barley agroinfiltration were transformed into *A. tumefaciens* COR308. Organelle markers with a mCherry or a GFP tag in plasmid pBIN20 [31] were transformed in *A. tumefaciens* LBA4404 or GV3101 respectively. P-bodies marker 35S:mCherry-DCP1 in vector pEAQ was transformed into *A. tumefaciens* GV3101.

For agroinfiltration assays, *A. tumefaciens* with the corresponding plasmid were grown at 28°C in Luria Bertani (LB) Miller or YM broth supplemented with appropriate antibiotics, 10 mM MES and 20 μM acetosyringone until an OD_600_ of 0.8-1. Cells were spun down at 4000 x g for 6 min at 4°C and then resuspended in fresh infiltration buffer (10 mM MES, 10 mM MgCl_2_ and 200 μM acetosyringone) to an OD_600_ of 0.3. The resuspended cells were then incubated at room temperature for a minimum of 3 hours. If appropriate, *A. tumefaciens* with different plasmids (e.g. organelle markers) were mixed 1:1 v/v before infiltration for a final OD_600_ of 0.3-0.6. A minimum of three *Nb* plants was agroinfiltrated with each construct or combination of constructs using a syringe into the leaves of 3-5 week-old plants. The same procedure was also used to agroinfiltrate 10-15 day-old leaves of cv. Odessa (1st and 2nd leaf) with target constructs along with a construct expressing RNA-silencing suppressor p19 at a 1:1 ratio to a final OD_600_ of 1.0.

### 2.5. Overexpression viral vector system (VOX)

The FoMV-based overexpression system (VOX) using PV101 derived vectors was used to infect *Nb* and barley plants as described in [21]. UhAvr1+SP, UhAvr1-SP and GFP were amplified by PCR using primers 2176 + 2178, 2177 + 2178 and 2174 + 2175, respectively, from previous entry vectors. All these PCR amplicons contained additional ClaI and XbaI sites for restriction cloning into vector pV101. The final VOX constructs, VOX:UhAvr1+SP, VOX:UhAvr1-SP and VOX:GFP, were verified by sequencing before transformation into *A. tumefaciens* GV3101 pSoup (Figure S1E). In each experimental repeat, a minimum of three *Nb* plants (3-4 week-old) were agroinfiltrated with each construct. The sap from agroinfiltrated leaves was used to rub-inoculate a minimum of eight barley plants (L1 and L2 leaf of 7 day-old plants).

### 2.6. Nb cell death assay

The GateWay™ LR reaction was performed between UhAvr1-SP entry vector and a modified pMCG161 vector (NCBI ID AY572837, ChromDB) to generate pMCG161:UhAvr1-SP:GFP and pMCG161:GFP:UhAvr1-SP (Figure S1F). As controls, empty pBIN61, pBIN61:GFP and pEAQ:GFP vectors [32,33] were used. Constructs were transformed into *A. tumefaciens* strain C58C1. For the assay, overnight cultures of *A. tumefaciens* were harvested at 4000g for 10 min at 4°C, resuspended in 10 mM MgCl_2_ to a final OD_600_ of 0.4 and infiltrated into the leaves of *Nb* plants using a syringe without needle. One day post-infiltration, cell death inducers (INF1, a PAMP-like elicitor from *P. infestans* [34], auto active resistance gene *Rx* [35,36] and a combination of *Bs2* a resistant gene from pepper and its cognate avirulence gene *AvrBs2* from *Xanthomonas campestris* pv. *vesicatoria* [37] were infiltrated in the same region as before at a final OD_600_ of 0.1. For *Bs2 and AvrBs2, A. tumefaciens* cultures were mixed 1:1 (v/v) for a final OD_600_ of 0.1. Symptoms were monitored up to 4 days after the second infiltration. A minimum of two plants (three leaves per plant) were infiltrated in each experiment.

### 2.7. PTI suppression assay

Constructs for specific use in *Psa* were based on GateWay™ destination vector pPSV_PsSPdes [38]. The pENTR/D-mCherry entry vector was constructed by PCR amplification of mCherry using primers 1504 + 1505 followed by GateWay™ TOPO cloning. The pENTR/D-mCherry and two previously generated entry clones (pENTR/D-UH_10022-SP and pENTR/D-UH_10022, [16]) were inserted in pPSV_PsSPdes using GateWay™ LR recombinase, generating PsSP:mCherry, PsSP:UhAvr1-SP and PsSP:UhAvr1+SP, respectively (Figure S1G). The *Pseudomonas* effector AvrRpm1 SP sequence, required for transfer of proteins into the host via the T3SS, was removed from pPSV_PsSPdes by a three piece ligation. The PCR amplification of a 139 bp DNA fragment upstream of AvrRpm1 SP was done using primers 1770 + 1777, and a 2.1 kb fragment downstream of AvrRpm1 SP was amplified using primers 1771 + 1776. These two fragments were then ligated with BglII-digested pVSP_PsSPdes, generating pVSPΔSPΔRFC-A. Then, mCherry or UhAvr1-SP was amplified by PCR using primers 1764 + 1765 and ligated into the EcoRV site of pVSPΔSPΔRFC-A resulting in ΔPsSP:UhAvr1-SP and ΔPsSP:mCherry (Figure S1H). Constructs were introduced into *Pseudomonas* following the procedure described in [39]. *Psa* cells cultured in liquid media were pelleted, washed twice with ice-cold 10 mM MgSO_4_ and resuspended in minimal medium (50 mM potassium phosphate buffer, 7.6 mM (NH_4_)2SO_4_, 1.7 mM MgCl_2_ and 1.7 mM NaCl, pH 5.7) with 10 mM fructose and appropriate antibiotics to an OD_600_ of 1.0. Then, cells were kept at 18°C overnight for induction, harvested at 4000g for 10 min at 4°C and resuspended in 10 mM MgCl_2_ to an OD_600_ of 1.0 for infiltration into the first leaves of 10-14 day-old plants of cv. Odessa. Leaf symptoms were scored at 48 hours post-inoculation (hpi).

Bacterial population density from infected plant tissue was measured as described in [40] with some modifications. Briefly, harvested leaves were surface sterilized in 70% EtOH, rinsed in sterile distilled water and dried on paper towels. The wide end of a 200 μl pipette tip was used to punch out leaf disks from four independent plants which were then pooled, representing one tissue sample. Pooled leaf disks were ground with water and 1:10 serial dilutions were made for each tissue sample followed by plating on LB agar plates containing appropriate antibiotics. Three such tissue samples were generated for each time point.

### 2.8. Barley protoplast transfection

Barley protoplasts were extracted from leaves of 6-10 day-old plants that were cut in slices with a sterile scalpel and digested in an enzyme solution (0.6 M Mannitol, 10 mM MES (pH 5.7), 1.5% Cellulase R-10 (Yakult Pharmaceutical, Japan), 0.75% Macerozyme R-10 (Yakult Pharmaceutical, Japan), 10 mM CaCl_2_ and 0.1% BSA) for 4-5 hours in the dark at room temperature with gentle shaking. Protoplasts were then overlaid with a salt solution (150 mM NaCl, 125 mM CaCl_2_, 5 mM KCl, 6 mM glucose, pH 5.8-6) followed by centrifugation at 250 x g for 10 min. Then they were washed with the salt solution, centrifuged for 5 min at 250g, resuspended in the salt solution again and incubated for 30 min at 4°C. Before transfection, the protoplasts were centrifuged at 250 x g for 5 min and resuspended in a transfection buffer (4 mM MES pH 5.7, 0.4 M mannitol and 125 mM MgCl_2_) at a final concentration of 10^6^-10^7^ protoplasts/ml. For each construct, 300 μl of protoplast suspension, 300 μl of PEG solution (40% PEG, 10% DMSO (dimethyl sulfoxide), 0.5 M mannitol, 0.1 M Ca(NO_3_)_2_, pH 7-9) and 15-30 μg of plasmid DNA were mixed gently. The mixture was incubated for 20-30 min in the dark and then mixed with 10 ml of minimal medium (0.7 M mannitol, 10 mM CaCl_2_, 1 mM KNO_3_, 1 mM MgSO_4_, 0.2 mM KH_2_PO_4_, pH 5.7). Then, the suspension was incubated at 4°Cfor 15 min, spun for 5 min at 250g, resuspended in 4 ml of minimal medium and kept in the dark at room temperature. Protoplasts were collected 24 hours post-transfection (hpt) by centrifugation at 1700g for 4 min.

### 2.9. Brefeldin A inhibitor assay

Brefeldin A (BFA) is a chemical inhibitor of the conventional ER-Golgi secretion pathway [41]. BFA (Invitrogen) assays in *Nb* were conducted on the 24 hours post-agroinfiltrated *Nb* leaves. Half of a leaf was infiltrated with water and the other half was infiltrated with 50 μg/ml BFA. These leaves were collected for protein blot analyses and confocal microscopy from three hours post-exposure. Several areas of a minimum of two leaves per construct were observed with confocal microscopy for each experimental repeat.

For BFA assays in fungal cells, 25-50 ml of cultured cells at an OD_600_ of 0.4-0.8 were spun down and resuspended in fresh medium, now supplemented with 50 μg/ml or 100 μg/ml of BFA or DMSO. The amount of DMSO corresponds to the amount used to make the BFA stock solution. The Uh1440 (Uh1351 (*MAT-1 UhAvr1* [*SP:UmPit2:mCherry*]) cells grown at higher OD_600_ of 0.6-0.8 died in the solution, therefore 25-80 ml of Uh1440 cells were grown to a lower OD_600_ of 0.2-0.6. After BFA exposure for 4 to 8 hours, the fungal cells were pelleted and their supernatants collected for tricarboxylic acid (TCA, Sigma-Aldrich) precipitation. Mean band intensities were calculated using Image Lab Software (Bio-Rad).

### 2.10. Pathogenicity assays

Haploid strains of *Uh* were grown until an OD_600_ of 1.0, mixed 1:1 v/v with Uh362 of opposite mating type and grown for an additional 24 hours to allow mating. The cultures were used to vacuum infiltrate the barley seeds as described in [16]. The percentage of infected plants was calculated by counting the total number of infected plants that showed at least one smutted head out of the total of inoculated plants. Total percentage of infected plants was calculated from two experimental repeats that consisted of a minimum of 15 plants each.

### 2.11. RNA extraction and quantification

Barley seeds were dehulled, surface sterilized and placed on sterile filter paper in Petri dishes to germinate in the dark at 22°C for 24 hours. Coleoptiles of the germinated seedlings were rub-inoculated using sterile cotton buds covered with teliospores, or soaked in water for the control set. Barley coleoptiles infected with teliospores were collected from 24 to 48 hpi from Petri dishes and pooled for RNA extraction. For 72 to 144 hpi samples, 48 hpi-infected seedlings were transplanted into potting mix and grown in a growth cabinet. Seedlings were rinsed with water, their crown tissue harvested, pooled and used for RNA extraction. Also, emerging barley heads were collected four months after germination and flash-frozen in liquid nitrogen until RNA extraction. Similarly, haploid fungal cells were harvested from liquid culture by spinning at 6000g for 15 min at 4°C and stored at −80°C until RNA extraction.

The samples were ground in liquid nitrogen and total RNA was extracted using a RNeasy Plant Mini Kit (Qiagen) along with on-column DNA digestion (RNase-Free DDase, Qiagen) for removal of contaminating genomic DNA. cDNA from the samples was synthesized using 0.5-1 μg of total RNA in a 20 μl reaction with Superscript VILO master mix (Invitrogen) following the manufacturers’ recommendation. The expression level of specific genes in these cDNA pools was quantified using a QX200 droplet digital PCR (ddPCR) system (BioRad) with gene-specific primers and QX200 EvaGreen Supermix (BioRad) following the manufacturer’s protocol. *UhAvr1* was detected by primer pair 1967 + 1968 that is specific to the region of *UhAvr1* and absent in the mutant Uh1289 (*MAT-1 ΔUhAvr1*). Primer pair 1965 + 1966 that targets *UhGAPDH* (*Uh glyceraldehyde 3-phosphate dehydrogenase*, NCBI gene ID DQ352820) was used to normalize the expression of each fungal gene of interest. In *Nb* agroinfiltrated samples, quantification of *UhAvr1* expression was done using the *UhAvr1* specific primers and their expression was normalized against the expression of the *Nb protein phosphatase 2* (*PP2A*) [42] as reference gene targeted by primer pair 2224 + 2225.

To calculate the fungal biomass in barley seedlings indirectly, the ratio between the expression of the fungal and the barley reference genes (*UhGapdh*/*HvUbiq*) was calculated. The expression of *HvUbiq* (NCBI Gene ID AK365157), a conserved homolog to *Triticum aestivum ubiquitin-conjugating enzyme* (NCBI gene ID AY736121) which was stably expressed in RNA-sequencing data from our lab and was found to be a proper reference gene in barley [43], was measured in the ddPCR assays using primer pair 520 + 521.

### 2.12. Extraction and detection of proteins

Leaves from a minimum of three barley or *Nb* plants were pooled and ground in protein extraction buffer (20 mM Tris pH 7.5, 5 mM MgCl_2_, 2.5 mM DTT, 300 mM NaCl, 0.1% NP-40, and ‘complete protease inhibitor cocktail’ (Roche)). Total protein lysates were collected after centrifugation at 3700g for 10 min at 4°C. Similarly, for protein extraction from fungal cells, fungal pellets were washed one time with 10 mM MgCl_2_ and ground in the presence of protein extraction buffer. To precipitate secreted protein from the spent liquid media, fungal cells were removed, then the supernatant was filter-sterilized through a 0.2 μm filter (VWR) to remove stray cells, and used for harvesting secreted proteins as in [44] with some modifications. Briefly, proteins from the supernatant were extracted by precipitating with 10% TCA overnight at 4°C followed by one cold acetate wash and resuspension in 2x loading buffer (50 mM Tris HCl pH 6.8, 2% SDS, 10% glycerol, 1% B-mercaptoethanol, 12.5 mM EDTA and 0.02% Bromophenol blue) before loading on acrylamide gels.

Immunoblots were performed as described in [16]. Briefly, proteins in loading buffer were boiled for 5 min before loading on to 12-15% SDS-PAGE gels using the Mini-Protean III gel system (BioRad). A protein transfer apparatus (BioRad) was used for transferring proteins to a Sequi-Blot PVDF membrane (BioRad) following the manufacturer’s protocol. Membranes were probed with commercial mouse monoclonal antibodies: anti-GFP (Clontech Living Colors JL-8), anti-mCherry (Abcam 1C51) or anti-tubulin (Calbiochem DM1A). UhAVR1 was detected using a custom made antirabbit polyclonal antibody (anti-UhAVR1, GenScript USA Inc). The peptide used to make this antibody is described in Table S4. Coomassie blue staining of the membranes was performed and used as loading control.

### 2.13. Staining and microscopy

The progression of infection by *Uh* teliospores on barley was visualized following a published protocol [45], with some modifications. A mixed staining solution consisting of 10 μg/ml of WGA-AF488 (Molecular Probes), 20 μg/ml of Propidium Iodide (PI; Sigma-Aldrich), 0.02% Tween 20 in phosphate-buffered saline (pH 7.4, 137 mM NaCl, 2.7 mM KCl, 8 mM Na_2_HPO_4_ and 1.5 mM KH_2_PO_4_), was used to vacuum infiltrate infected barley seedlings for 30 min. Then, the samples were washed twice in phosphate-buffered saline and kept in this solution until visualization. The fungal dye, Uvitex-2B (Polysciences Inc., USA), was occasionally used in combination with PI as described in [46] with the following modifications: infected barley seedlings were stained in 0.1% w/v Uvitex 2B in 0.1M Tris-HCl (pH 5.8) for 15 min. Then, seedlings were subjected to a few washes with water followed by staining with 10 μg/ml of PI for 15 min. Samples were washed again a few times with water before visualization. Stained coleoptiles and first leaves were longitudinally dissected before placement on a microscope slide. The same procedure except staining was performed for tracking the GFP fluorescence *in planta* upon infection with teliospores.

Confocal microscopy was performed with a Leica TCS SP8 confocal laser scanning microscope with a 63x water immersion objective in a sequential scanning mode. Each experiment consisted of a minimum of two slides originating from independent samples (e.g. leaf tissue, seedlings). In each slide, a minimum of three spots (e.g. cells) were visualized. During the visualization, WGA-AF 488 was detected by excitation at 490 nm and emission at 500-530 nm. PI was detected by excitation at 561 nm and emission at 591-651 nm. Uvitex 2B was detected by excitation at 405 nm and emission at 410-440 nm. GFP was detected by excitation at 480/488 nm and emission at 495-550 nm. mCherry was detected by excitation at 587 nm and emission at 598-644 nm. Chlorophyll was detected by excitation at 480 nm and emission at 680-730 nm. The percentage of PI stained hyphae on infected barley seedlings at 48 hpi was calculated by counting the number of confocal images with at least a PI stained hypha (each image represents an infection site) out of the total of images in all experimental repeats performed.

## 3. Results

### 3.1. Fungal growth is affected by the deletion of UhAvr1 at the early stage of barley infection

Previously, the expression of *UhAvr1* was detected at 48 hpi on infected seedlings of resistant cv. Hannchen leading to localized cell death and complete immunity [2,16]. To further elucidate the function of *UhAvr1*, a detailed time-course infection using *Uh* teliospores with or without *UhAvr1* was carried out on susceptible cultivar Odessa (*ruh1*) and resistant cultivar Hannchen (*Ruh1*). To resemble the natural infection process, either the wild type (WT, resulting from a cross of Uh362 x Uh364) or UhAvr1m (*UhAvr1* deletion mutant, Uh362 x Uh1289) teliospores were applied on the surface of barley coleoptiles (for strain numbers and genotypes, see Table S1). At 24 hpi, similar fungal germination and growth from both types of teliospores on both cultivars were observed (Figure 1). Sporidial mating on the coleoptile surfaces of both cultivars infected with either WT or UhAvr1m teliospores was visible at this time point (Figure 1, first row).

**Figure 1.**
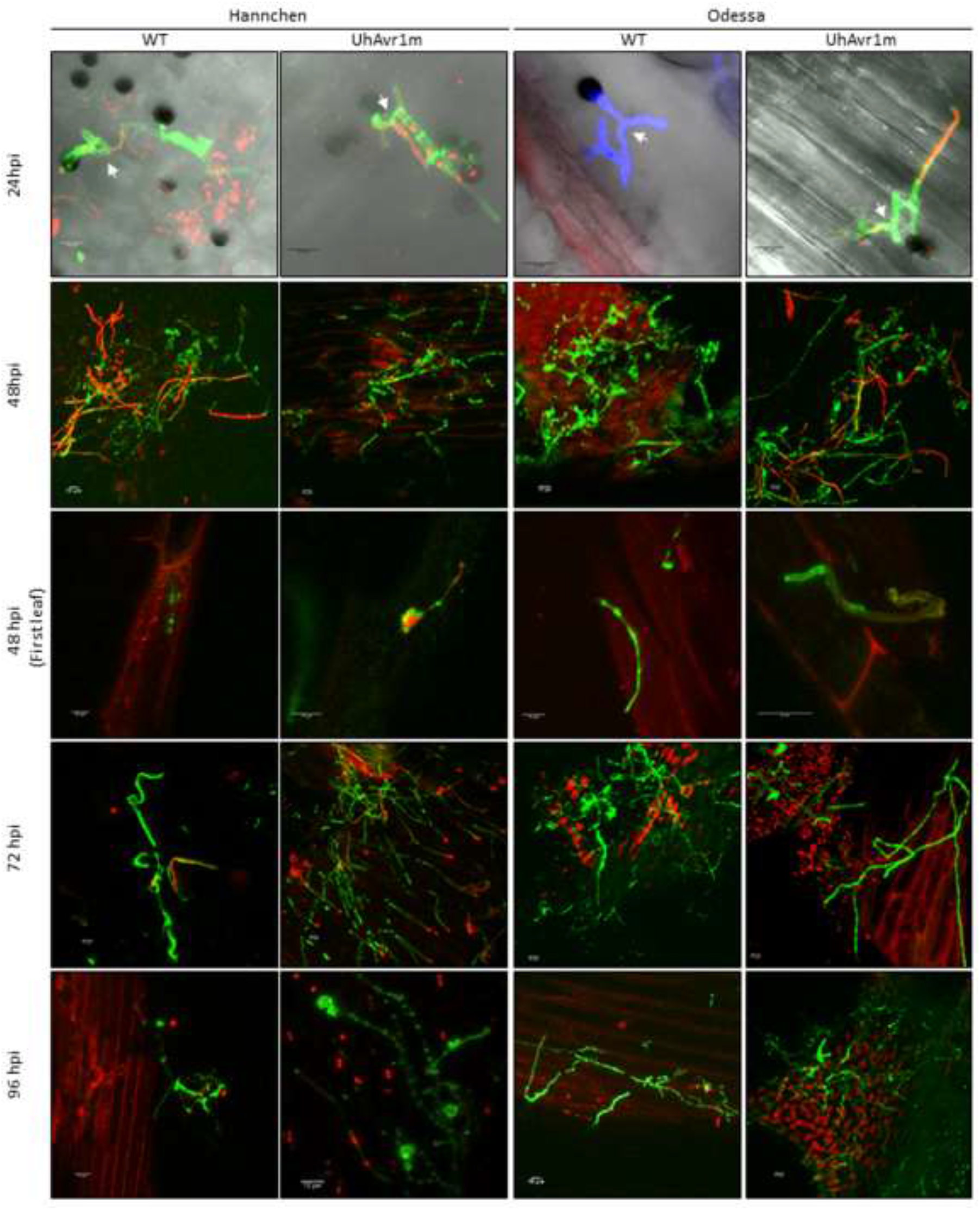
Progression of infection by *U. hordei* during early stages. Confocal microscopy of barley cv. Odessa (*ruh1*) and Hannchen (*Ruh1*) shows differential fungal growth and infection by WT or UhAvr1m teliospores at early stages of infection. Tissues were stained with WGA-AF488 (green) or Uvitex2B (blue) together with PI (red). Teliospore germination, sporidia formation and mating via conjugation hyphae (arrow) were seen on the coleoptile surfaces of both cultivars at 24 hpi. At 48 hpi, dead hyphae (red) were observed which were prominent for WT teliospores on cv. Hannchen and for UhAvr1m teliospores on cv. Odessa. Lower fungal growth from UhAvr1m teliospores at 72 hpi on the cv. Odessa recovered by 96 hpi. Similarly, reduced fungal growth from WT teliospores was seen in the cv. Hannchen starting from 48 hpi. Scale bar represents 10 μm. All images are maximum intensity projections of confocal stacks except for 24 hpi Odessa WT which is a confocal snapshot of a single optical section. Confocal imaging was performed on a minimum of three infected seedlings. Experiments were repeated two times with similar results and representative images are displayed

At 48 hpi, dead hyphae stained with Propidium Iodide (PI) were seen on cv. Hannchen infected with WT teliospores (~70% of visualized sites) but to a lesser extent on plants infected with UhAvr1m teliospores (~40% of visualized sites) suggesting plant defence triggering at this stage (Figure 1, second row). At this point, no PI-stained barley cells could be found on the coleoptile but fungal hyphae were observed on the first leaf of plants infected with both teliospores (Figure 1, third row) suggesting penetration had already occurred at this stage. At 72 hpi and onwards, there was extensive fungal growth on coleoptiles inoculated with UhAvr1m teliospores compared to the fungal growth on coleoptiles inoculated with WT teliospores (Figure 1, fourth and fifth row). This is consistent with the fact that *UhAvr1* triggers plant defence leading to fungal death in the incompatible interaction [2,4,16].

On the cv. Odessa at 48 hpi, dead hyphae stained with PI were mostly observed on the surface of coleoptiles inoculated with UhAvr1m teliospores (~85% of visualized sites) compared to coleoptiles inoculated with WT teliospores (~40% of visualized sites), suggesting suppression of plant defence by *UhAvr1* (Figure 1, second row). Also, similar to the infection on the cv. Hannchen, WT and UhAvr1m fungal hyphae were found on the first leaf of cv. Odessa indicating fungal penetration prior to this time point (Figure 1, third row). Previously, penetration of fungal hyphae during the compatible interaction was shown to be initiated from 30 hpi onwards, followed by inter- and intracellular fungal growth [4]. At 72 hpi, fewer fungal hyphae were observed on coleoptiles inoculated with UhAvr1m teliospores compared to coleoptiles inoculated with WT teliospores, indicating a drop in vigour upon deletion of this effector (Figure 1, fourth row). At later stages, fungal growth from UhAvr1m teliospores recovered as seen by extensive hyphal growth on the coleoptiles (Figure 1, fifth row). This suggests that on a compatible host, UhAVR1 plays a role in optimizing early infection, possibly suppressing defence responses until other effectors come to play a role at later stages.

Many plant pathogenic fungi develop an appressorium that is crucial for host tissue penetration and infection [47,48]. An undifferentiated appressorium structure characterized by a hyphal tip swelling has been reported in *Um* [49,50] and a similar appressorium-like structure was also observed for *Uh* [3]. These structures from *Uh* and *Um* were seen in the juncture between two adjacent cells [3,50]. Knob-like fungal structures, that under confocal microscopy appear different from the previously described appressorium-like structures, were observed at 48 hpi onwards on the coleoptiles of both cultivars infected with both types of teliospores but not at all infection sites (Figure 2). These structures were observed mainly on or nearby the junction between two neighbouring barley cells (Figure 2 and Figure S2), though, it was technically difficult to track the origin of these structures due to the collapse of older hyphae (Figure S2). Further characterization of this structure and the thin septated hyphae emanating from it, could not be conducted with confocal microscopy due to the thickness and rigidity of the coleoptile tissue. However, we hypothesize that this structure is an appressorium-like structure which aids *Uh* during the infection process.

**Figure 2.**
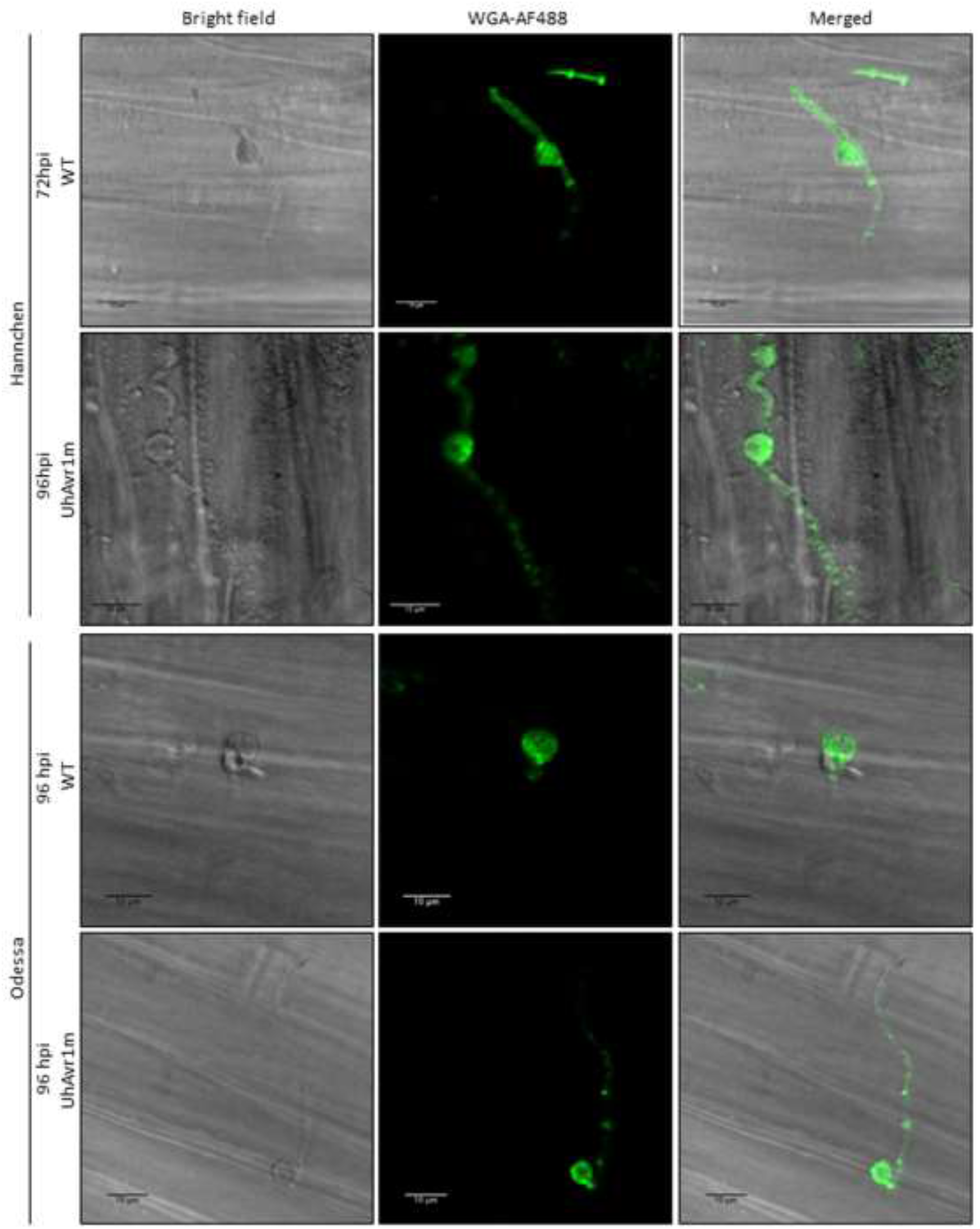
Appressorium-like structures are seen in both types of interaction. Seedlings from the cv. Odessa and cv. Hannchen were infected with WT or UhAvr1m teliospores followed by staining with WGA-AF488 (green). A putative appressorium with a knob-like structure was seen during imaging of coleoptiles 48 hpi. Thin septated hyphae emanating from the appressorium-like structure are noticeable in confocal microscopy that differ from the non-septated hyphae found anterior to it. Scale bar represents 10 μm. All images are a snapshot of a single z-stack.

### 3.2. UhAvr1 is expressed in planta during early infection of barley seedlings

Towards elucidating the function of this effector, a detailed understanding of its expression profile at different infection stages is needed. For this, *UhAvr1* transcript levels from infected barley seedlings and liquid cultured haploid cells were measured. The assay was specific for *UhAvr1* since no transcripts were detected in the mock control and in both barley cultivars infected with UhAvr1m teliospores. Very low expression of this effector was detected in liquid cultured haploid Uh364 cells (*MAT-1, UhAvr1*) but higher expression of *UhAvr1* was detected on the host infected with WT teliospores. Steady increases of *UhAvr1* transcripts were detected from 24 to 96 hpi in WT teliospores while infecting both cultivars (Figure 3A). However, the levels of *UhAvr1* transcript decreased in cv. Odessa while transcripts were not detectable in cv. Hannchen at 144 hpi. Interestingly, very low expression of this effector was detected in smutted barley heads (Figure 3A). Overall, the pattern of expression suggests that the functionality of this effector is time-specific, particularly during early stages of infection.

**Figure 3.**
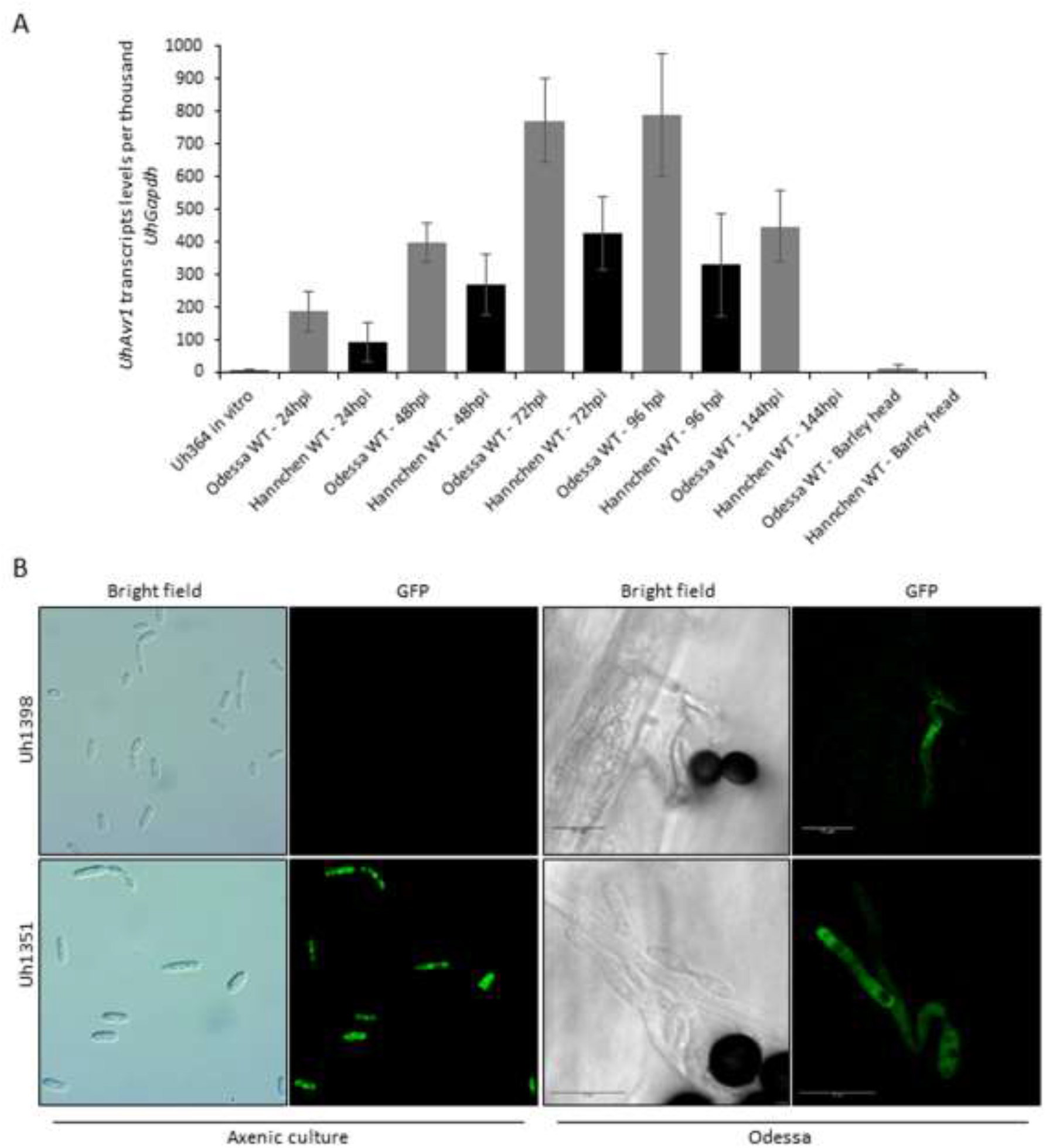
*UhAvr1* expresses at the early stage of infection. **(A)** Expression of *UhAvr1* in haploid cells and during the course of barley infection. Accumulation of *UhAvr1* transcripts was measured in haploid cells grown in liquid medium, in seedlings inoculated with WT teliospores, and in heads of cv. Hannchen and cv. Odessa. The graph shows the average from three independent biological repeats each consisting of three technical repeats for ddPCR. Values are expressed in number of *UhAvr1* transcripts per 1000 reference gene *UhGAPDH* transcripts. Error bars represent the standard deviations calculated from the three biological repeats. **(B)** Microscopy images showing UhAVR1 expression upon host sensing. Haploid cells of strain Uh1351, constitutively expressing GFP, showed fluorescence while haploid strain Uh1398, expressing *UhAvr1* tagged GFP from its WT promoter (SP:GFP:UhAvr1), did not. Infection of cv. Odessa with teliospores from cross Uh362 x Uh1351, as well as SGUhAvr1 teliospores (from cross Uh362 x Uh1398), showed GFP fluorescence at 40 hpi. Microscopy of haploid strains grown in liquid media was done four times independently (Axioscope, Carl Zeiss); in barley, two independent repeats were performed. All confocal images are a snapshot from a single optical section and the scale bar represents 10 μm.

To verify this result and correlate with protein production, we modified the mutant strain Uh1289 by replacing *ΔUhAvr1* with *SP:GFP:UhAvr1* expressed from its wild type *UhAvr1* chromosomal location and promoter, generating a new haploid strain, Uh1398. As a control, we used a previously generated strain Uh1351 [16] that expresses GFP from a strong constitutive promoter (otef:GFP). Microscopy performed on fungal haploid cells grown in liquid media did not show any GFP fluorescence for Uh1398 but strong fluorescence for Uh1351 (Figure 3B). To track the expression of *UhAvr1* during plant infection on cv. Odessa, SGUhAvr1 (Signal Peptide:GFP:UhAvr1) teliospores were generated from a cross of Uh362 x Uh1398, and control GFP teliospores from a cross of Uh362 x Uh1351. Interestingly, GFP fluorescence emanating from the hyphae of SGUhAvr1 teliospores was also observed on the infected barley coleoptile, supporting the idea that *UhAvr1* is only turned on upon sensing the host (Figure 3B).

### 3.3. UhAVR1 counteracts early host defence responses and triggers an HR in Ruh1-harboring barley plants

In certain pathosystems, deletion of a fungal effector can interfere with the fungal growth, its ability to colonize its host and disease severity [51]. Previously, pathogenicity assays showed that the deletion of *UhAvr1* did not lead to a reduction of virulence in the susceptible cultivar [16]. However, the pathogenicity assays which assessed the number of plants showing infected seed heads at 2 to 3 months post-inoculation compared to the total number of inoculated plants, are likely not sensitive enough to detect subtle effects. Therefore, to monitor the effect on fungal growth during early infection caused by *UhAvr1*, we adopted an indirect approach by measuring changes in fungal biomass based on the expression of housekeeping genes of the host and the fungus. Calculations of the ratio of the expression of these genes at different time points showed an increase of the fungal biomass on cv. Hannchen upon inoculation with both WT and UhAvr1m teliospores at 24 and 48 hpi (Figure 4A). This corroborates the microscopic observations of germinating teliospores and resulting hyphal growth on the surface of the coleoptiles as seen in Figure 1. After 48 hpi, the deduced fungal biomass of the WT infection started to steadily decrease whereas the deduced fungal biomass resulting from UhAvr1m teliospores increased further (Figure 4A). This is in accordance with the higher number of PI-stained hyphae from WT teliospores observed on cv. Hannchen at 48 hpi (Figure 1), suggesting fungal death due to UhAVR1-triggered plant defence initiation.

**Figure 4.**
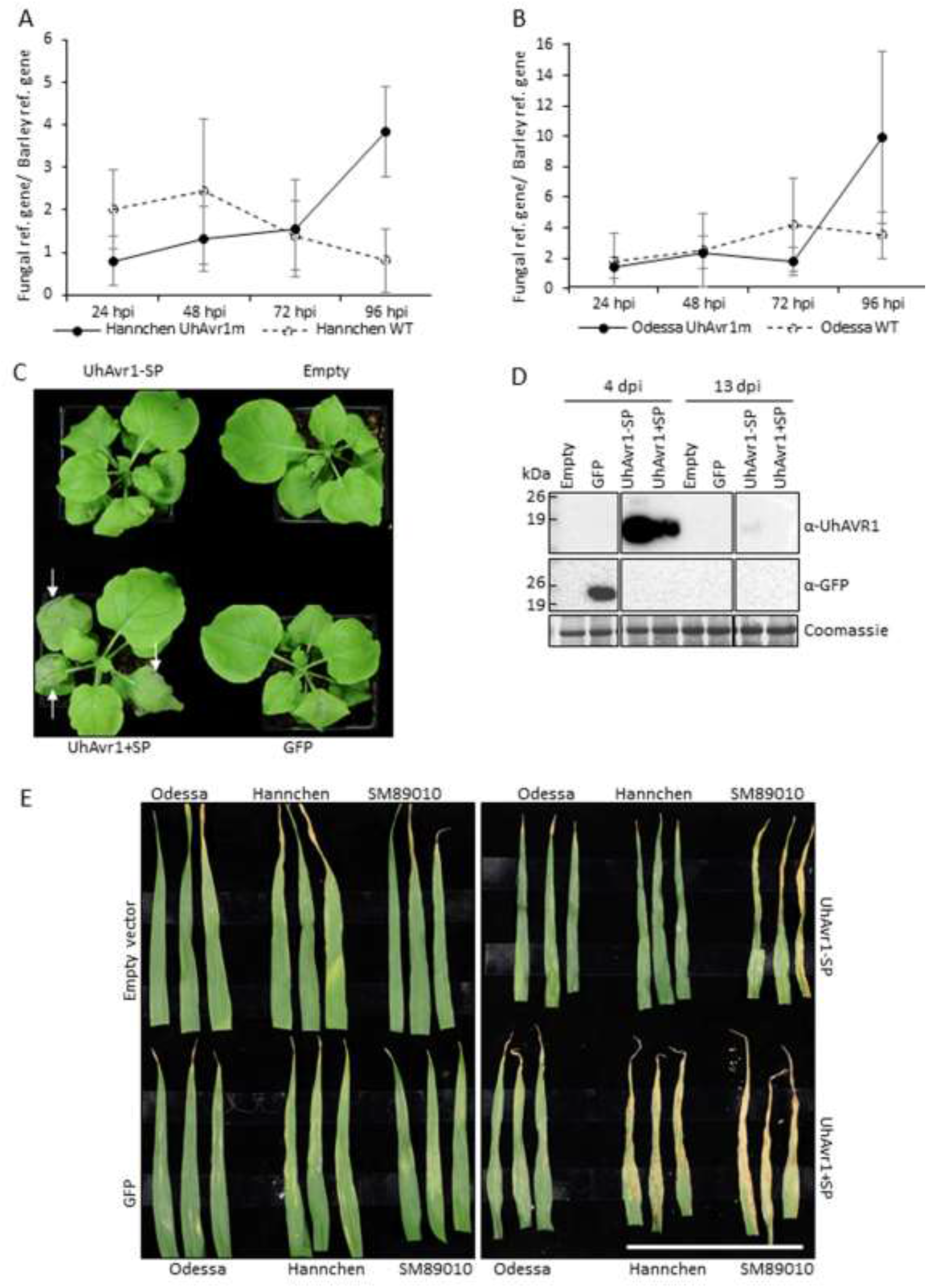
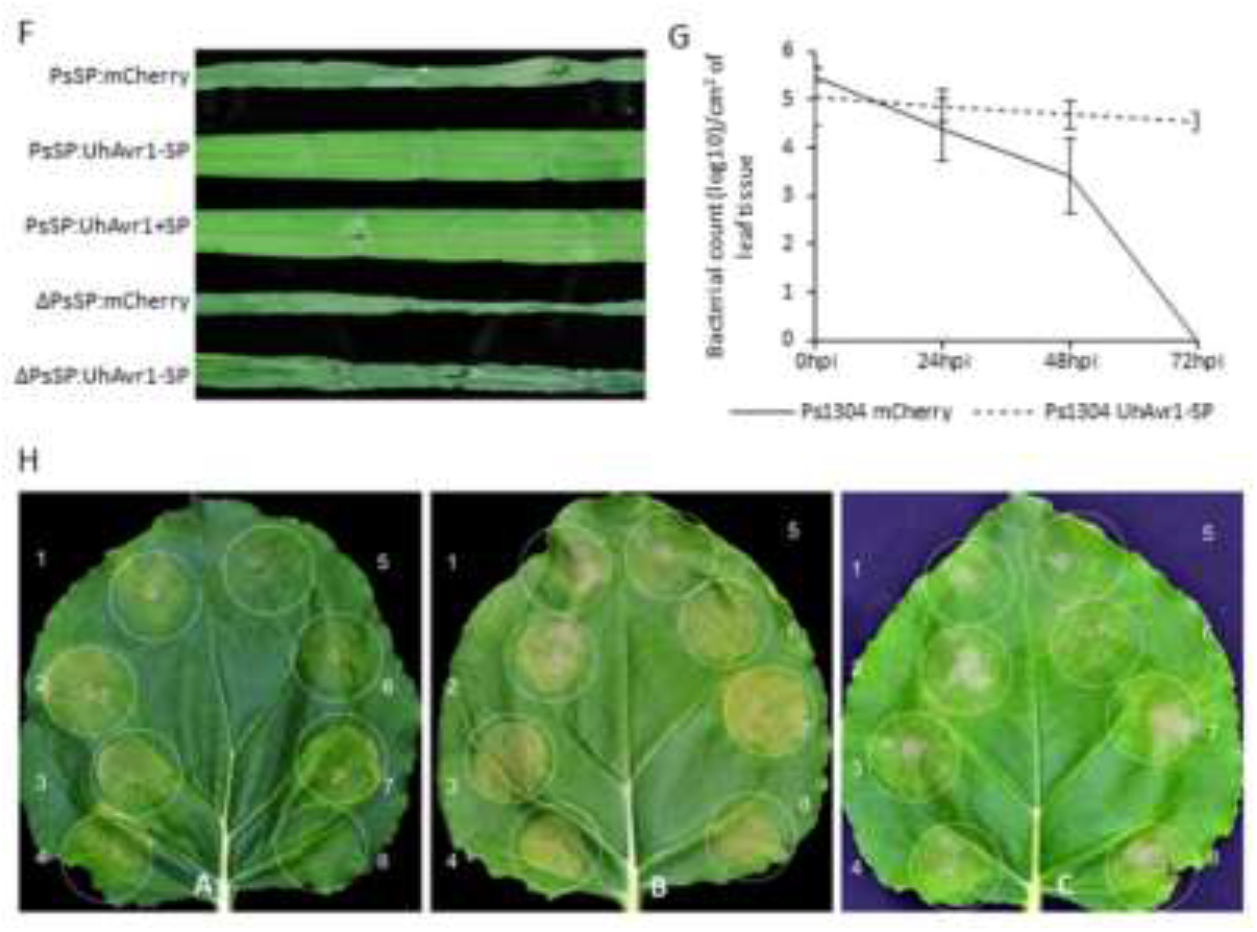
UhAVR1 triggers defence during the incompatible interaction and suppresses a conserved target of plant immunity during the compatible interaction. **(A)** Deduced fungal biomass during infection by WT or UhAvr1m teliospores in cv. Hannchen and **(B)** cv. Odessa. Fungal biomass was deduced by calculating the ratio between the expression levels of fungal (*UhGAPDH*) and barley (*HvUbiq*) reference transcripts. Differential fungal biomass was observed starting from 48 hpi. Graphs show the average of three independent biological repeats with error bars depicting the corresponding standard deviations. **(C)** *Nb* leaves agroinfiltrated with VOX:UhAvr1+SP triggered accelerated strong necrosis at 6 dpi (arrows) which was not yet observed with constructs expressing UhAVR1 lacking a SP, GFP and the empty vector. The experiment was repeated three times with similar results and a representatives image is shown. **(D)** Protein blots confirmed expression of UhAVR1 in *Nb* using the VOX system. Samples were collected from agroinfiltrated leaves at 4 dpi and upper non-inoculated leaves at 13 dpi from the same batch of plants shown in the image of panel C. The blots show the presence of the full-length UhAVR1-SP (19 kDa), UhAVR1+SP (21 kDa) and GFP (27 kDa) at 4 dpi. At 13 dpi, UhAVR1-SP could be detected in the upper non-inoculated leaves. Coomassie staining of the large subunit of Rubisco is shown as loading control. Protein blot analysis performed on one of the experimental repeats is shown to verify virus infection and protein accumulation. The picture shown is cropped from the same blot membrane. **(E)** Resistant barley cultivars display HR-like symptoms upon VOX-mediated delivery of UhAvr1+SP. L2 leaves expressing VOX:UhAvr1+SP (white bar) exhibit strong HR-like symptoms in cvs. Hannchen and SM89010 carrying *Ruh1* at 13 dpi. Images show three representative L2 leaves from three different barley plants for each construct. All images were taken from the same experimental repeat using the same settings. **(F)** UhAVR1 suppresses *Pseudomonas*-induced cell death. mCherry, UhAvr1-SP and UhAvr1+SP with a bacterial T3SS SP sequence (PsSP) were delivered into the leaves of 10 day-old cv. Odessa using *P. syringae* pv. *atropurpurea* 1304 (*Psa* 1304). At 48 hpi, PsSP:mCherry triggered cell death but not the constructs PsSP:UhAvr1-SP or PsSP:UhAvr1+SP. Whereas, deletion of the *Pseudomonas* SP (ΔPsSP) could not prevent cell death for ΔPsSP:UhAvr1-SP. **(G)** *Psa* 1304 expressing UhAVR1-SP counteracts the nonhost resistance response in barley. Leaves of cv. Odessa were infiltrated with *Psa* 1304 expressing PsSP:mCherry or PsSP:UhAvr1-SP and bacterial counts were made up to 72 hpi. The bacterial population of *Psa* 1304 expressing PsSP:mCherry declined drastically (six-fold) whereas the population expressing PsSP:UhAvr1-SP declined only slightly. The graph shows the average of a minimum of three experimental repeats for each time point with their corresponding standard deviation shown as error bars. **(H)** UhAVR1 suppresses plant cell death induced by three elicitors in *Nb*. *Nb* leaves were agroinfiltrated with: (1) empty pMCG161:GW:GFP, (2) empty pMCG161:GFP:GW, (3) pEAQ:GFP, (4) pBIN61:GFP, (5) pMCG161:UhAvr1-SP:GFP, (6) pMCG161:GFP:UhAvr1-SP, (7) pEAQ:GFP or (8) pBIN61:GFP. Twenty-four hours later, the same regions were agroinfiltrated with three different cell death inducers: (A) *AvrBs2/Bs2*, (B) auto-active RX or (C) INF1. UhAvr1-SP:GFP (5) and GFP:UhAvr1-SP (6) suppressed cell death induced by all three elicitors. Pictures were taken 4 days post-inoculation of the cell death inducers.

On cv. Odessa, deduced fungal biomass increased at 24 and 48 hpi on plants inoculated with both types of teliospores, similar to on cv. Hannchen (Figure 4B). However, at 72 hpi, there was a decrease in deduced fungal biomass for UhAvr1m compared with the WT teliospore infection (Figure 4B) coinciding with the high levels of PI-stained hyphae at 48 hpi on coleoptiles of cv. Odessa inoculated with UhAvr1m teliospores (Figure 1). Although a better resolution of fungal growth inside the plant could not be achieved due to technical difficulties (e.g. very low amount of fungal tissue in comparison to the plant tissue), results from both microscopy and deduced fungal biomass measurements converge on the idea that *UhAvr1* is a virulence factor and its function is related to suppressing plant defence responses during the initial stage of barley infection.

To further test the hypothesis that this effector may be suppressing plant defence responses, a virus-mediated overexpression vector (VOX) system based on FoMV was used [21]. VOX constructs (VOX:UhAvr1+SP, VOX:UhAvr1-SP, VOX:GFP or VOX:empty) were first agroinfiltrated in *Nb* leaves to establish viral infection. Sap from *Nb* leaves was then further used as inoculum to infect one week-old barley leaves of cv. Odessa (*ruh1*), cv. Hannchen (*Ruh1*) and cv. SM89010 (*Ruh1*) [20]. We observed significant differences in the timing of appearance of necrosis in agroinfiltrated *Nb* leaves depending on the constructs used. When infiltrating VOX:UhAvr1+SP, necrosis appeared at 3 dpi which had further accelerated by 6-8 dpi, whereas weak necrosis appeared for the same batches of plants agroinfiltrated with VOX:UhAvr1-SP or VOX:GFP from 6-8 dpi (Figure 4C). By 13 dpi, leaves agroinfiltrated with all four constructs showed necrosis (Figure S3A). Protein blot analyses confirmed the expression of the UhAVR1-SP and UhAVR1+SP proteins in the agroinfiltrated *Nb* leaves at 4 dpi and expression from UhAVR1-SP in the systemically infected leaves at 13 dpi (Figure 4D). Accumulation of UhAVR1 protein expressed from the UhAvr1+SP construct was lower than expression from UhAvr1-SP construct at 4 dpi and could not be detected at 13 dpi which was expected as the necrosis appeared earlier on UhAvr1+SP infiltrated leaves. Accelerated necrosis caused by the expression of UhAvr1+SP in *Nb* suggests its role in increasing susceptibility to either the virus or to the *Agrobacterium* strain used and hence supporting a role in suppressing (general) defence responses.

The resistant barley cultivars inoculated with sap derived from *Nb* agroinfiltrated with VOX:UhAvr1+SP showed HR-like symptoms consisting of necrotic streaks starting from 7 dpi in the L2 leaves. These HR-like symptoms were stronger in both resistant cultivars (cv. Hannchen and SM89010) at 13 dpi (Figure 4E). Whereas, L2 leaves of cv. Odessa inoculated with *Nb*-derived VOX:UhAvr1+SP lacked any of these symptoms at 13 dpi. Also, HR-like symptoms on the L2 leaves were absent for all three cultivars inoculated with the other constructs (VOX:empty, VOX:GFP and VOX:UhAvr1-SP) up to 15 dpi (Figure 4E). These results show that VOX-mediated expression of *UhAvr1* in barley leads to HR-like symptoms in the cultivars carrying *Ruh1*. Occasionally, VOX:UhAvr1+SP inoculated plants of cv. Odessa showed systemic mosaic symptoms on the upper L3 leaves at around 15 dpi (1-2 plants out of 8 plants inoculated in each of the repeats; Figure S3B). Although the frequency of mosaic viral symptoms was low, this result supports the idea that this effector is enhancing the susceptibility to FoMV in cv. Odessa. Overall, the VOX-based experiments in these two plant systems suggest that UhAVR1 suppresses a conserved component(s) of plant immunity leading to increased susceptibility to other pathogens (virus and/or *Agrobacterium*) in both barley and *Nb*.

To further investigate the role of UhAVR1 in suppressing general immunity of plants, *Pseudomonas*-based assays were conducted on the barley cv. Odessa. UhAvr1-SP, UhAvr1+SP and mCherry were delivered into the leaves of 10 day-old barley via the Type III Secretion System (T3SS), using the vector-based *P. syringae*-specific signal peptide (PsSP). The *P. syringae* pv. *atropurpurea* strain 1304 (*Psa* 1304) used in this study, causes a nonhost reaction (cell death) in barley. At 48 hpi, cell death was observed in plants infiltrated with *Psa* carrying mCherry whereas no visible symptoms were observed in plants infiltrated with *Psa* harbouring UhAvr1-SP or UhAvr1+SP, suggesting UhAVR1 suppressed the nonhost reaction triggered by *Psa* (Figure 4F). To confirm that this effect was due to the action of UhAVR1 secreted by the T3SS, a construct was generated in which the *Pseudomonas* secretion SP was deleted (ΔPsSP). Without the PsSP, cell death was seen in plants infiltrated with *Psa* carrying mCherry but now also for UhAvr1-SP at 48 hpi (Figure 4F). In support, bacterial counts in leaves of cv. Odessa infiltrated with a suspension of 1 x 10^6^ CFU/ml induced bacteria harbouring constructs PsSP:mCherry or PsSP:UhAvr1-SP, showed that PsSP:mCherry populations in the leaf tissue decreased six-fold within three days following inoculation, whereas PsSP:UhAvr1-SP populations declined only slightly (Figure 4G). The reduction in the bacterial population expressing PsSP:mCherry supports an *in planta* suppression triggered by a nonhost reaction against this bacterium whereas a limited reduction of the PsSP:UhAvr1-SP expressing population supports a role for UhAVR1 in Pattern-Triggered Immunity (PTI) suppression.

To further refine the role of UhAVR1 in suppressing immunity-associated cell death, three different types of cell death inducers, *INF1, Rx* and a combination of *Bs2* with *AvrBs2*, were tested in *Nb*. The *INF1* elicitor from *P. infestans* is suggested to be delivered into the apoplast by this oomycete and is later on internalized into the cytoplasm of the host cells where it interacts with a cellular protein triggering defence responses in *Nb* [52]. *Rx* is a nucleocytoplasmic CC-NBS-LRR resistance gene against Potato Virus X (PVX) which triggers an HR response upon recognition of the viral coat protein in *Nicotiana* [53]. *AvrBs2* is an avirulence effector from *Xanthomonas campestris* pv. *vesicatoria* which is secreted by the T3SS and translocated into host cells. Pepper plants harbouring *Bs2* recognize this effector leading to cell death and plant immunity [54]. As expected, cell death in leaves infiltrated with elicitors in combination with control vectors (empty or GFP) was observed (Figure 4H). Interestingly, both N- or C-terminally GFP-tagged UhAVR1 chimeric proteins (GFP:UhAVR1-SP and UhAVR1-SP:GFP) suppressed tissue necrosis generated by all three cell death inducers but at different degrees. This suppression seemed more pronounced when using the C-terminal GFP-tagged UhAVR1 construct in combination with all three elicitors (Figure 4H). This set of experiments suggests that UhAVR1 suppresses cell death in this heterologous system and triggered by various pathways. Also, it suggests that UhAVR1 virulence function is not affected by the GFP tag which is not the case for its avirulence function in barley.

### 3.4. The signal peptide of UhAVR1 is essential for secretion and is sensitive to Brefeldin A

Confocal microscopy to localize endogenously expressed SP:GFP:UhAVR1 (using strain Uh1398) was challenging due to low levels of expression. Thus, a new fungal secretion system (FunGus) to express and deliver effector proteins was created (Figure 5A). A haploid strain (Uh1430) expressing UhAVR1+SP:mCherry from an episomal plasmid and constitutively expressing genome-integrated GFP was generated. The liquid grown Uh1430 exhibited mCherry fluorescence, indicating proper expression of UhAVR1+SP:mCherry from the episomal plasmid (Figure 5B 1-3). Mating of this strain with Uh362 (*Uhavr1*) of opposite mating type was not affected as seen in the typical ‘fuzz’ reaction on charcoal plates and mated cells in microscopy (Figure 5B 4-6 and 5C). Also, its conjugating hyphae exhibited mCherry fluorescence suggesting the effector was moving along with the fungal cytosol (Figure 5B 4-6). Proteins extracted from pelleted Uh1430 cells (intra-cellular proteins, P) as well as the supernatant (secreted proteins, S) showed the presence of chimeric UhAVR1 (Figure 5D). We observed the SP-cleaved UhAVR1:mCherry along with some breakdown products. As a control, an antibody against tubulin was used to validate the absence of cell contents in the liquid media (Figure 5D) confirming the secretion of chimeric UhAVR1 from intact cells. To confirm that the SP was responsible for the secretion of this effector, an *Uh* strain Uh1434 was generated, where the SP of UhAVR1 was deleted (pUHES:ΔSP:UhAvr1-SP:mCherry). Proteins collected from the pellet of Uh1434 cells but not from the supernatant showed UhAVR1 (Figure 5D). Detection of UhAVR1 in the supernatant of Uh1430 but not of Uh1434 showed that the deletion of the SP is enough to block the secretion of UhAVR1 into the liquid media and potentially into the host.

**Figure 5.**
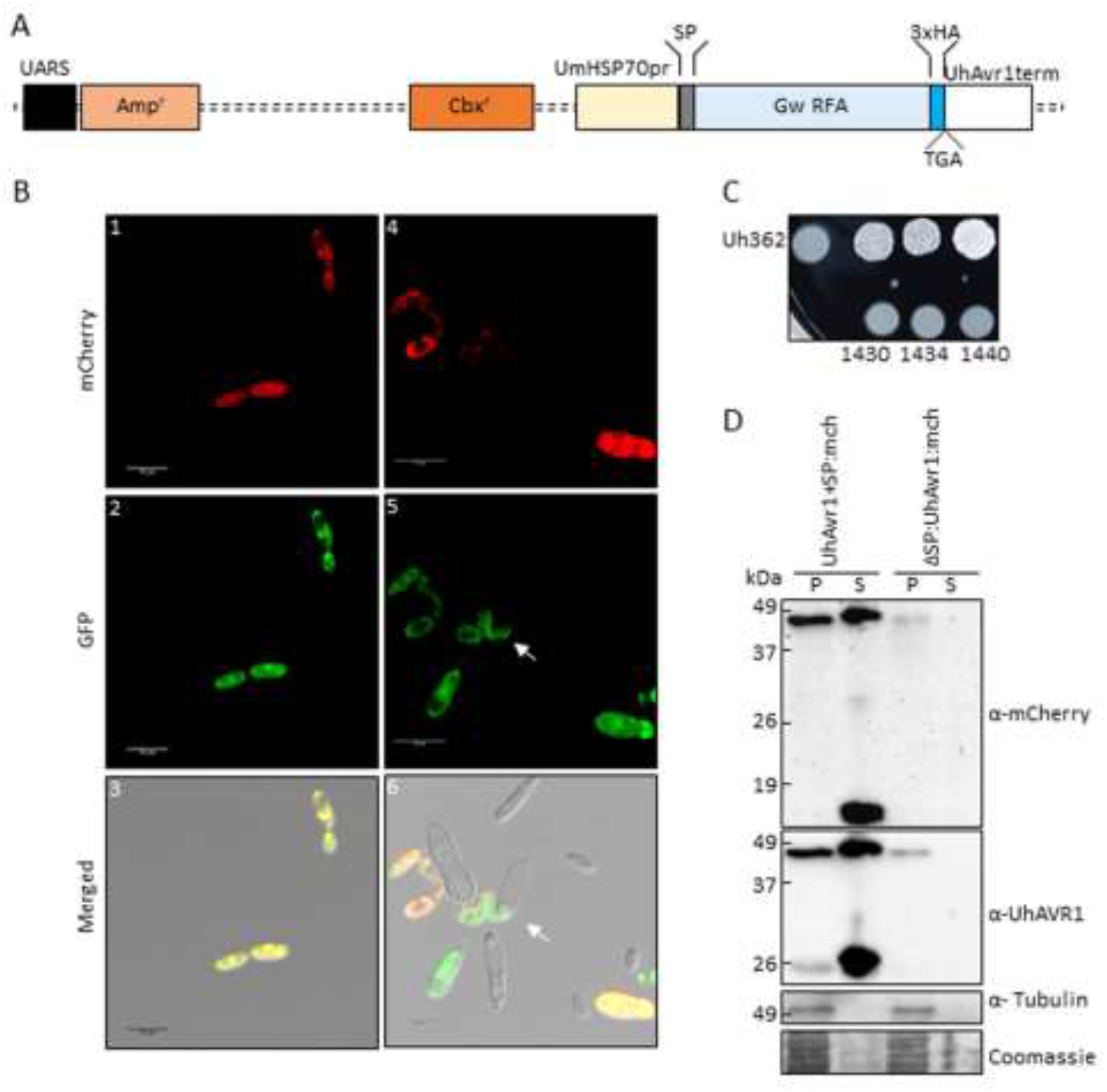
A new *Ustilago* secretion system (FunGus) efficiently expresses and secretes fungal effector UhAVR1. **(A)** Graphical representation of FunGus constructs. GateWay™ (Invitrogen) based vector (pUHESdest) possesses a *Ustilago* Autonomously Replicating Sequence (UARS), bacterial (Amp^r^) and *Ustilago* (Cbx^r^) antibiotic resistance genes for selection. Effector proteins inserted via GateWay™ recombineering will be expressed under the control of the *Um* HSP70 constitutive promoter (*UmHSP70pr*) with the N-terminal UhAVR1 SP. The presence of a 3xHA tag followed by a stop codon (TGA) provides the choice of adding this epitope to the effectors. **(B)** Confocal microscopy of *Uh* expressing UhAVR1:mCherry from the FunGus system: **(1-3)** haploid strain Uh1430 grown in liquid media with mCherry fluorescence derived from episomal UhAvr1+SP:mCherry and GFP fluorescence derived from genome-integrated transgene otef:GFP. **(4-6)** in mated cells (Uh1430 x Uh362), mCherry fluorescence from Uh1430 moves together with cytosolic GFP fluorescence into conjugation hyphae (arrows). A snapshot of a single optical section is displayed for all confocal images and the scale bar represents 10 μm. **(C)** Three independent *Uh* transformants expressing FunGus-based cassettes (Uh1430, Uh1434 and Uh1440) are not affected in mating showing a typical “fuzzy” phenotype on a charcoal plate. **(D)** Deletion of the UhAVR1 SP in the FunGus cassette blocks secretion of the effector into the liquid medium. Protein blots of Uh1430 (UhAvr1+SP:mCherry) but not of Uh1434 (ΔSP:UhAvr1-SP:mCherry) showed SP-cleaved UhAVR1:mCherry in the spent medium (secreted proteins, S) though the protein is detected in the pelleted cells (intracellular proteins, P) in both strains. Proteins observed in the spent medium are not due to cell lysis, depicted by the lack of Tubulin (60 kDa) in the medium. The predicted sizes of UhAvr1+SP:mCherry, UhAvr1-SP:mCherry and ΔSP:UhAvr1:mCherry are 50, 46 and 46 kDa, respectively. Confocal imaging of Uh1430 haploid cells in (B) and protein blot in (C) are from the same set of samples. Protein blots of Uh1430 and Uh1434 cells were performed three times with similar results.

This finding suggests a conventional pathway of secretion via the ER-Golgi. However, recent studies in *Phytophthora infestans* [55] have shown that apoplastic and cytoplasmic effectors are delivered by different secretion pathways. To further investigate the possible pathway of secretion of UhAVR1, we performed comparative experiments also using a known apoplastic effector with cysteine proteases inhibitor activity from *Um*: *UmPit2* [56,57]. We constructed a strain (Uh1440) expressing UmPit2+SP:mCherry, having its natural SP, from an episomal construct while also constitutively expressing GFP. Uh1430 and Uh1440 were grown in liquid medium and exposed to BFA or DMSO as a control. The SP-cleaved UhAVR1:mCherry along with breakdown products were detected in the pelleted cells that had undergone either treatment. However, lower amounts of UhAVR1:mCherry protein were detected in the spent medium upon BFA exposure (Figure 6).

Similarly, for UmPit2:mCherry, the predicted full-length protein as well as the SP-cleaved protein were detected in the pelleted cells upon either treatment (Figure 6). Also here, the accumulation of cleaved product for UmPit2:mCherry was less in spent medium in samples treated with BFA compared with DMSO (Figure 6). Unlike UmPit2:mCherry, UhAVR1:mCherry was present in only a single protein band in the spent medium and pelleted cells, though the apparent molecular weight of these two bands was different (Figure 6). However, both proteins were likely of the same size but migrated slightly differently in the gel due to differences in the isolation procedure; fungal cell pellets generated higher amounts of more complex proteins and cellular debris than the proteins extracted from the spent media. Also, proteins were precipitated from the spent medium with TCA, and traces of TCA could have affected the pH and hence mobility on gels. No intracellular protein, tubulin, could be detected in the spent medium of all samples when probed with anti-tubulin antibody ruling out leakage or lysis of fungal cells (Figure 6). These experiments were challenging to perform as higher concentrations of or longer exposure to BFA led to fungal cell death; this was ruled out in the presented experiment by probing blots with an internal control (anti-tubulin). Altogether, these results show that secretion of UhAVR1 and UmPit2 is sensitive to BFA confirming they are routed via the conventional ER-Golgi pathway.

**Figure 6.**
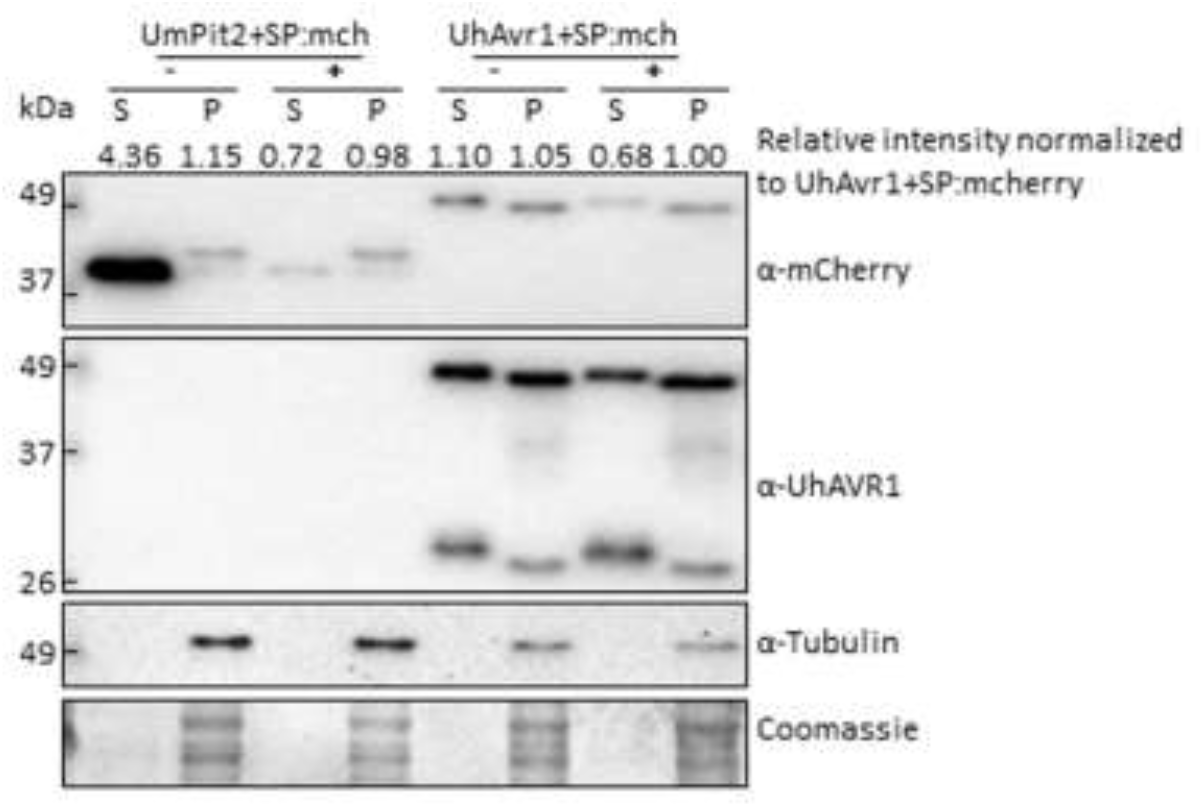
Brefeldin A exposure affects the secretion of UhAVR1 and UmPit2 to liquid medium. Cells of Uh1430 (UhAvr1+SP:mCherry) and Uh1440 (UmPit2+SP:mCherry) were grown in liquid medium and exposed to BFA 100μg/ml (+) or DMSO (-) for 4 hours. Protein blot analysis of proteins extracted from pelleted cells (intracellular proteins, P) and from spent medium (S) revealed that both effectors were secreted into the medium with secretion reduced upon exposure to BFA (+). The predicted sizes are: UmPit2+SP:mCherry (40 kDa), UmPit2-SP:mCherry (37 kDa), UhAvr1+SP:mCherry (50 kDa) and UhAvr1-SP:mCherry (46 kDa). Experiments were performed three times independently for UmPit2+SP:mCherry and five times for UhAvr1+SP:mCherry with similar results. The intensity value of each protein band detected by anti-mCherry antibody was calculated by Image Lab software and provided on the top of the gel picture.

### 3.5. UhAVR1 localizes to the cytosol of plant cells

Attempts to track and localize this effector *in planta* using the FunGus secretion system were not successful possibly due to the high expression of chimeric UhAVR1 and its instability (Figure 5D) affecting visualization by microscopy. Instead, SGUhAvr1 (Uh362 x Uh1398) teliospores with lower expression of UhAVR1 from the natural *UhAvr1* promoter were used to track SP:GFP:UhAVR1. This is a similar system to the described FunGus system as it relies on the fungal secretion system to secrete UhAVR1. Teliospore-infected cv. Odessa showed GFP fluorescence in between barley cells that remained intact even after plasmolysis of the tissue (Figure S4A and 4B). Similar results were obtained for the UhAvr1GFP teliospores (Uh362 x Uh1357) expressing C-terminally GFP-tagged UhAVR1 from the strong constitutive *otef* promoter (Figure S4C and S4D). This is likely the site of fungal secretion of UhAVR1 which is later presumably taken up by the cell. However, many attempts to confirm the stability of chimeric proteins in barley by protein blots were unsuccessful probably due to the low number of infection sites leading to low levels of UhAVR1 protein and due to the detection of unspecific barley proteins by the anti-UhAVR1 antibody.

To understand if these chimeric proteins are properly directed to their target site and functioning, we performed pathogenicity assays using mated strains expressing SP:GFP:UhAVR1 (Uh1397, Uh1398 and Uh1399). Loss of avirulence on cv. Hannchen compared to infection with mated WT strains (Uh362 x Uh364) was seen for these strains, likely due to the interference of the N-terminal fluorescent moiety on the HR-triggering function of UhAVR1 (Table S5). Previous work also showed a loss of avirulence upon infection of cv. Hannchen with C-terminally GFP tagged UhAVR1 strains expressed from the *UhAvr1* wild type promoter (*UhAvr1:GFP*, Uh1353) as well as from the constitutive *otef* promoter (*otef*:*UhAvr1:GFP*, Uh1357) [16]. In that study we also showed that smaller and charged tags such as HA added to the C-terminus of UhAVR1 led to the loss of avirulence [16]. Our results show that both N-terminal and C-terminal extensions on the UhAVR1 protein are interfering with its avirulence function and could interfere with its translocation to its target location for function in barley. Similarly, translocation assays done in maize infected with *Um* strains expressing fluorescently tagged effectors failed to observe translocation of effectors into plant cells [58].

In light of this, we decided to use protoplasts from cv. Hannchen or cv. Odessa transfected with similar gene constructs, now expressed from the plant-specific 35S promoter, 35S:GFP, 35S:UhAvr1-SP:GFP or 35S:UhAvr1+SP:GFP, to verify whether such chimers would be detectable in the cytoplasm. Protoplasts of both cultivars displayed GFP fluorescence in the cytosol and occasionally in cytosolic foci for both UhAvr1-SP:GFP and UhAvr1+SP:GFP constructs (Figure 7, second and third row). These foci occasionally appeared in the 35S:GFP control only when a stronger GFP signal was obtained, suggesting they could be artefacts of over-expression (Figure 7, first row). Protein blot analyses of these transfected protoplasts could not detect UhAVR1 because of very low levels of expression and hence the stability of the chimers could not be ascertained.

**Figure 7.**
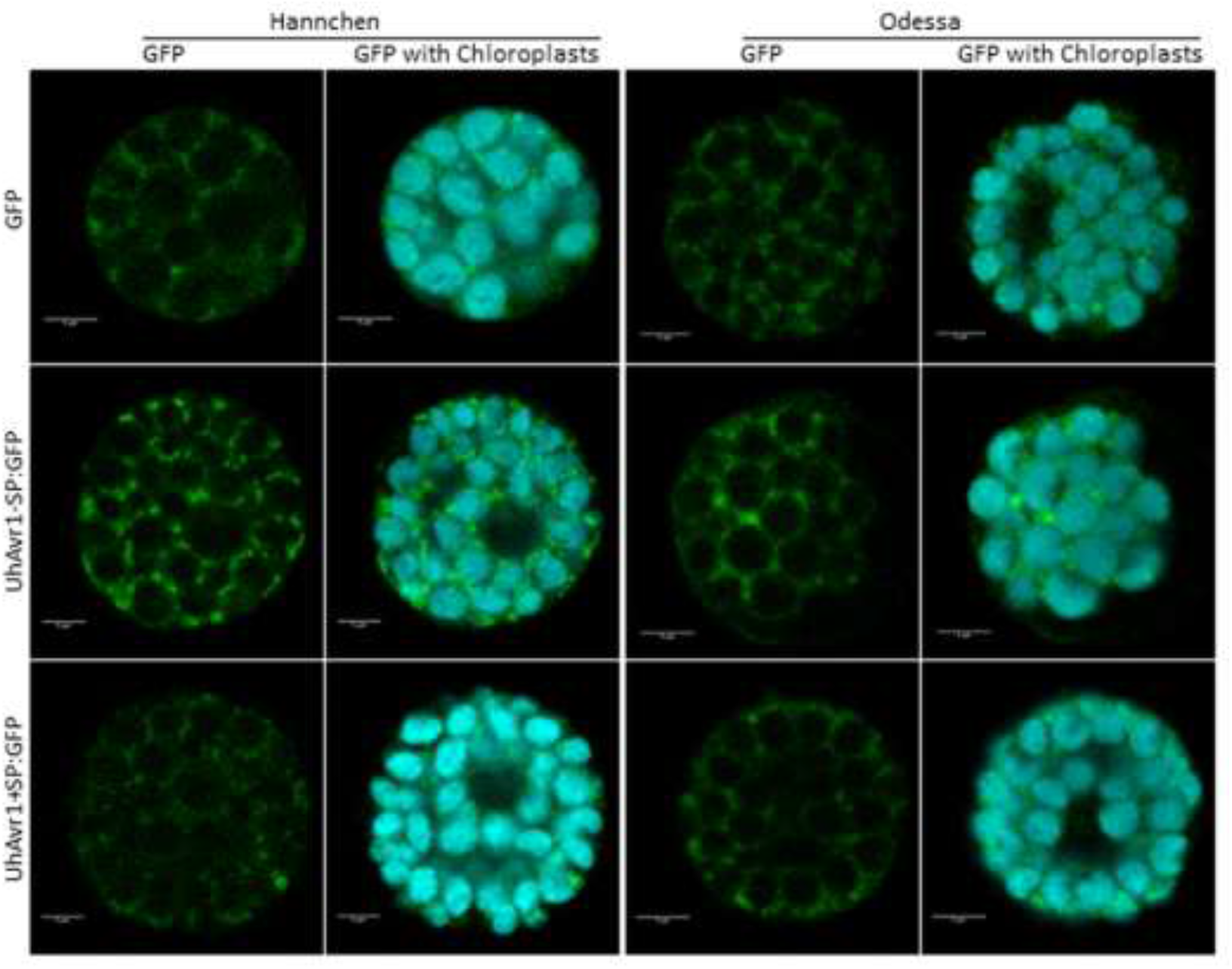
UhAVR1 localizes to the cytosol of barley protoplasts. Confocal imaging of protoplasts of cv. Odessa and cv. Hannchen 20 hrs post-transfection with 35S:GFP, 35S:UhAvr1-SP:GFP and 35S:UhAvr1+SP:GFP. GFP fluorescence for all transiently expressed constructs in both cultivars localized to the cytosol surrounding chloroplasts and occasionally in cytosolic foci. Three experimental repeats were performed for cv. Odessa and two for cv. Hannchen with similar results. Scale bar for confocal images represents 5 μm. A snapshot of a single optical section is displayed for all images.

Protoplasts cannot provide an indication as to whether fungal secreted UhAVR1:GFP chimers are being targeted to the cytoplasm, unlike expression in tissue with possible delivery into the apoplast. To further elucidate localization of this effector, we tested the same constructs but now delivered by agroinfiltration into leaves of 10-15 day-old barley cv. Odessa. A similar cytosolic localization was seen in both UhAvr1-SP:GFP and UhAvr1+SP:GFP infiltrated leaves validating our observation made in protoplasts. The free GFP control showed nucleoplasm and cytosolic localization (Figure S5). The efficiency of *Agrobacterium* delivery, expression and the resulting intensity of the fluorescence was low in barley plants, making this approach challenging and preventing proper visualization on protein blots.

The study of UhAVR1 in its natural host presented many challenges such as low expression levels, unstable chimeric protein and the inability to detect UhAVR1 from barley samples on protein blots. To circumvent these problems, the same constructs used for barley were used for transient expression in a heterologous system, *Nb*, via agroinfiltration. At 24 hpi, GFP fluorescence was observed in the nucleoplasm and the cytosol of the agroinfiltrated plants expressing free GFP (Figure 8A). Similarly, in the UhAvr1-SP:GFP agroinfiltrated plants, fluorescence was observed in the nucleoplasm, cytosol and in some occasional cytosolic foci of different sizes (Figure 8A). Whereas for the UhAvr1+SP:GFP agroinfiltrated plants, fluorescence was observed only in the cytosol and cytosolic foci of various sizes were always present, while occasional protein aggregation in the cytosol was seen at 24 hpi (Figure 8A). Expressing UhAvr1+SP:GFP (49 kDa) resulted in lower GFP fluorescence and protein accumulation on protein blots than when agroinfiltrating and expressing

**Figure 8.**
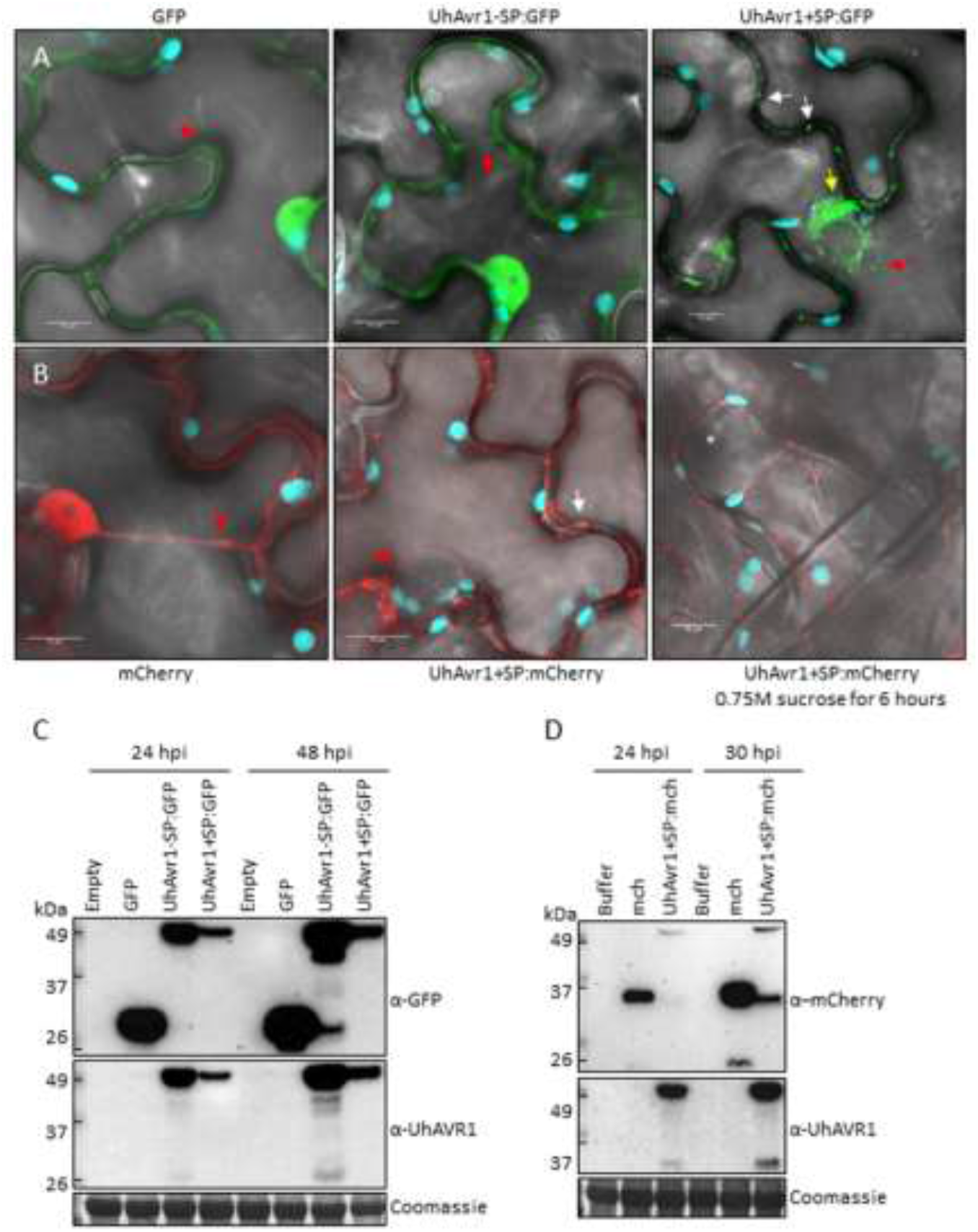
UhAVR1 localizes to the cytosol of *Nb* cells. Confocal imaging of *Nb* leaves 24 hours after agroinfiltration with constructs expressing fluorescent UhAVR1 chimers. **(A)** Free GFP (control) and UhAvr1-SP:GFP fluorescence localized to the nucleoplasm and cytosol (red arrow) whereas UhAvr1+SP:GFP fluorescence localized to the cytosol (red arrow) and cytosolic foci (white arrows). Protein aggregation in the cytosol could be occasionally seen at 24 hpi (yellow arrow). **(B)** Fluorescence from free mCherry was seen in the nucleoplasm and cytosol (red arrow) whereas, when infiltrating construct UhAvr1+SP:mCherry, fluorescence was seen in the cytosol (red arrow) and cytosolic foci (white arrow) at 24 hpi. Cells expressing UhAvr1+SP:mCherry were subjected to plasmolysis with 0.75M sucrose for 6 hours; no mCherry could be detected in the apoplastic space (asterisk). **(C)** Protein blots of the same batch of plants used for panel A showed the expression of intact GFP, UhAvr1-SP:GFP (47 kDa) and UhAvr1+SP:GFP (49 kDa) at 24 and 48 hpi. **(D)** Protein blots from the same batch of plants used for panel B showed mCherry (27 kDa) and UhAVR1:mCherry (+SP of 48 kDa and −SP of 46 kDa) at 24 hpi and 30 hpi (plasmolysis). Three independent repeats of confocal microscopy and protein blots were performed and representative results are shown. Images of panel A and B were produced by merging GFP or mCherry, chlorophyll and bright field channels. A snapshot of a single optical section is displayed for all confocal images and scale bars represents 10 μm.

UhAvr1-SP:GFP (47 kDa) at 24 and 48 hpi despite these constructs being on the same vector backbone (Figure 8C). Therefore, we quantified transcription levels in these samples using ddPCR and showed less accumulation of *UhAvr1* transcripts for UhAvr1+SP:GFP compared to UhAvr1-SP:GFP, ~60% less at 24 hpi and ~30% less at 48 hpi (Figure S6A). However, the corresponding protein levels did not correlate well with the measured transcript levels as the accumulation of protein resulting from agro-infiltration of UhAvr1+SP:GFP was almost five times less at 24 hpi than that resulting from UhAvr1-SP:GFP; at 48 hpi it was almost one order of magnitude less (Figure S6B). Since the only difference in the transcripts is the 57 nt sequence coding for the SP, differential transcription or translation is unlikely to cause these discrepancies. Protein blot analysis showed that the products resulting from agroinfiltration of UhAvr1+SP:GFP or UhAvr1-SP:GFP were identical in size (Figure S6C) indicating processing when using the UhAvr1+SP:GFP construct to yield cleaved, mature UhAVR1-SP:GFP and hence transitioning through the ER-Golgi. Though the difference in protein levels could be due to protein instability in the ER/Golgi or derived vesicles, it is more likely that the protein is secreted from the host cell and is unstable in the apoplast.

We verified the cytosolic localization using mCherry as the fluorescent moiety by agroinfiltrating construct 35S:UhAvr1+SP:mCherry into *Nb*. The free mCherry control showed fluorescence in the nucleoplasm and cytosol of *Nb* leaf cells at 24hpi (Figure 8B). Whereas, UhAvr1+SP:mCherry fluorescence was seen only in the cytosol and in cytosolic foci of various sizes, though overall fluorescence was less than for the control (Figure 8B). The localization was similar to what was seen when using construct UhAvr1+SP:GFP. Plasmolysis was performed in cells expressing UhAvr1+SP:mCherry but no mCherry fluorescence was found in the apoplastic space at 30 hpi (Figure 8B). Protein blot analyses from these plants at 24 hpi and 30 hpi showed the presence of mCherry (27 kDa) and SP-cleaved UhAVR1:mCherry (46 kDa), indicating mostly intact chimers (Figure 8D).

The presence of cytosolic foci observed in *Nb* cells could be artefacts due to overexpression or may indeed be caused by the UhAVR1 localization to or in a cellular organelle. Therefore, 35S:UhAvr1+SP:GFP was co-expressed with several organelle markers in *Nb* (peroxisomes, Golgi, ER and P-bodies) or the agroinfiltrated cells were stained with PI (cell wall staining). The UhAVR1:GFP fluorescence did not localize with any of the organelle markers tested (Figure S7). This suggests that the visualized cytosolic foci may be an artefact of overexpression or represent unknown cell organelles.

### 3.6. The UhAVR1 fungal SP directs linked proteins through the Nb secretory pathway via the ER-Golgi

Since we could not detect UhAVR1 tagged with either GFP or mCherry in the apoplast, we questioned whether the SP of UhAVR1 can direct secretion out of *Nb* cells and whether re-entry occurs. Could it direct other proteins to the apoplast? To test this, the SP of UhAVR1 (19 aa plus an additional 6 aa after the predicted SP cleavage site) was attached to the N-terminus of the GFP protein generating 35S:SP:GFP. Confocal microscopy of 35S:SP:GFP expressing cells showed GFP fluorescence in the cytosol and cytosolic foci of different sizes (Figure S8A), similar to what was seen for UhAvr1+SP:GFP and UhAvr1+SP:mCherry (Figure 8A and 8B). Also, some *Nb* cells occasionally showed GFP protein aggregation in the cytosol. No apoplastic localization of GFP was found upon plasmolysis of cells (Figure S8B). Protein blot analysis of these plants showed two protein bands for SP:GFP suggesting cleavage of the SP near or at the junction with GFP by the host (Figure S8C). Due to the possibility of the GFP not fluorescing in the apoplastic space [59], we continued the experiments using mCherry-tagged constructs.

SP-mediated export of the protein to the apoplastic space may require some cellular factor(s) that could be absent in *Nb*. To answer this question, the functionally characterized *Um* apoplastic effector *UmPit2* was tagged with mCherry together with its original signal peptide (35S:UmPit2+SP:mCherry). Confocal microscopy at 24 hpi of *Nb* cells expressing UmPit2+SP:mCherry and a GFP-tagged plasma membrane marker showed apoplastic localization of mCherry as observed in between the plasma membrane marker (Figure S9A). Protein blot analysis of these plants revealed the presence of the SP-cleaved UmPit2:mCherry (Figure S9B). Overall, this shows that the UmPit2 SP is able to direct UmPit2 protein into the apoplast. Furthermore, it also shows the ability of a smut SP to work in a heterologous system suggesting the conservation of the pathway in both fungi [56] and *Nb*.

This led us to investigate whether the UhAVR1 SP was mediating the delivery of UhAVR1 protein expressed in *Nb* through the plant secretory pathway. *Nb* plants transiently expressing test constructs were exposed to BFA or water, for 3 hours. Confocal microscopy upon exposure to BFA showed the presence of protein aggregation in the cytosol in the *Nb* cells expressing constructs that possess a SP (SP:GFP, UhAvr1+SP:GFP and UhAvr1+SP:mCherry; Figure S10A) compared to the water control. Whereas, no difference was observed between BFA or water exposed cells when using constructs without SP (GFP and UhAvr1-SP:GFP) (Figure S10).

Exposure to BFA leads to fusion of the Golgi-ER and blocking of the transport from the ER to Golgi [41]. To verify if the protein aggregated in BFA exposed cells co-localized to the fused ER-Golgi, chimeric effectors 35S:Avr1+SP:mCherry or 35S:UmPit2+SP:mCherry were co-expressed with GFP-tagged Golgi and ER markers. Confocal imaging showed co-localization of UhAvr1+SP:mCherry and UmPit2+SP:mCherry with both markers in BFA exposed cells but not in water treated cells (Figure 9A). Protein blot analyses of these plants showed the presence of the SP-cleaved UhAVR1:mCherry and UmPit2:mCherry (Figure 9B). Also, breakdown chimeric mCherry products were detected for both effectors but the amount of these products was higher for both effectors upon water exposure compared with BFA exposure. This suggests secretion to the apoplast which leads to more protein degradation. Overall, these results support the narrative that the presence of the SP in these effectors leads them to enter and cross the secretory pathway of *Nb* through the ER-Golgi. Arguments for likely re-entry of UhAVR1-SP in to plant cells are presented in the Discussion.

**Figure 9.**
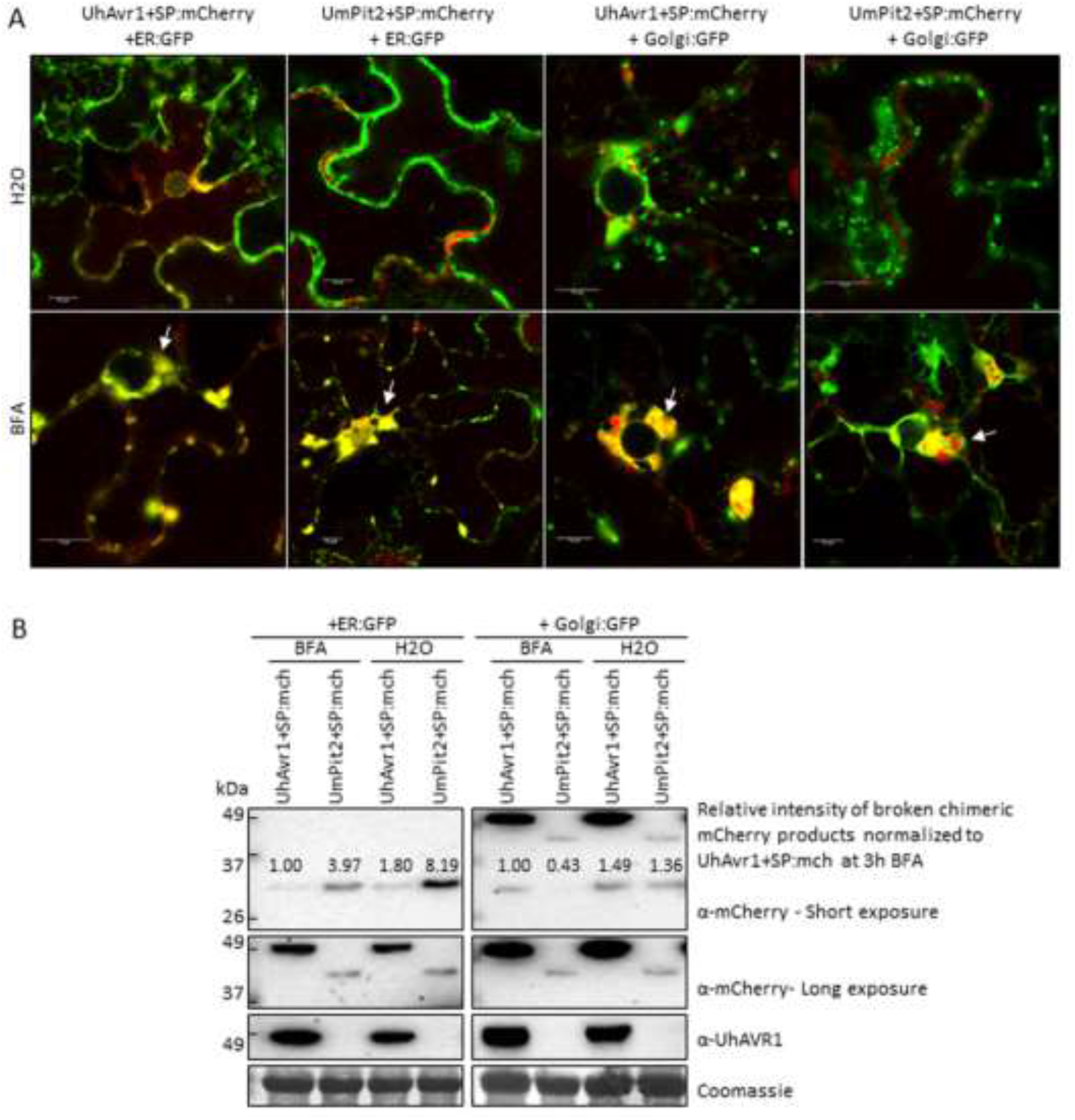
UhAVR1:mCherry and UmPit2:mCherry co-localize with fused ER-Golgi in *Nb* leaf cells upon BFA exposure. **A)** Confocal microscopy of mCherry-tagged fungal effectors which accumulate in secretory pathway organelles of *Nb* leaf cells upon BFA exposure. A GFP-tagged ER or Golgi organelle marker were co-agroinfiltrated with UhAvr1+SP:mCherry or UmPit2+SP:mCherry and exposed to BFA or water. mCherry fluorescence from both chimeric effectors (arrows) co-localized with the fused ER-Golgi (GFP fluorescence) after 4 hours of BFA exposure. UmPit2:mCherry and UhAVR1:mCherry fluorescence localized to the apoplast and cytosol, respectively, upon water exposure. Snapshots of a single optical section are displayed and the scale bar represents 10 μm. **(B)** Protein blot analyses from the same batch of plants used in panel A showed the SP-cleaved UhAVR1:mCherry and UmPit2:mCherry after 3 hours of BFA or water exposure. Breakdown products were detected only with anti-mCherry antibody. Immuno-blots were imaged for shorter or longer exposure to reveal unsaturated mCherry breakdown products and the full-length proteins. The intensity value of each breakdown chimeric mCherry product was calculated by Image Lab software and provided on the gel picture. Short exposure of blots probed with anti-mCherry antibody corresponds to 16 sec for the +ER:GFP and 2002 sec +Golgi:GFP. Whereas, long exposure of the same blots corresponds to 4429 sec for +ER:GFP and 2541 sec for +Golgi:GFP. Three independent repeats were done and a representative image is shown.

## 4. Discussion

In this study, we show that UhAVR1, the only avirulence effector identified in smuts to date, has a SP that is essential for secretion from fungal cells, a process that is inhibited by BFA and hence occurs via the ER-Golgi pathway. *UhAvr1* expression in *Uh* hyphae begins upon sensing of barley host cells and persists mainly during the initial stages of infection. During these first days, UhAVR1 has a role in virulence as it suppresses general defence responses including nonhost/PTI and ETI invoked by various pathogens (*Uh*, *Pseudomonas* spp., FoMV) in various plants (barley and *Nb*), indicating its target could be conserved among various plants. Barley therefore likely evolved to guard this target since interaction or action of UhAVR1 in cultivars having *Ruh1* triggers an HR, as we show using the FoMV system for effector delivery and expression. Finally, we show that UhAVR1 is found in the cytoplasm of plant cells, including in barley, suggesting this is the site of its action. We provide indirect evidence that mature UhAVR1, with its SP-cleaved off, can enter barley plant cells without the presence of *Uh* fungus.

### 4.1. Expression and function of UhAVR1

A time-course study using microscopy and ddPCR showed that *UhAvr1* expression is stage- and time-specific (Figure 1, 3A and 3B). Transcriptome analysis of leaves of susceptible barley (cv. Golden Promise) infected by *Uh* showed that 65% of all *in planta* induced *Uh* effector genes are stage-specific including *UhAvr1* [18], similar to our study for *UhAvr1* in cvs. Odessa and Hannchen. UhAVR1 lacks homology to any known protein or protein motifs making it difficult to deduce its function. Certain barley cultivars have evolved to possess a single dominant resistant gene, *Ruh1*, guarding against this effector or its action. Orthologues of *UhAvr1* can be found in other pathogenic smuts and in a fungus closely related to the smuts [17] suggesting its role as a core effector. Core effectors are generally conserved in related species and are hypothesized to enable early host interactions by overcoming plant immunity components (Zuo, Ökmen et al. 2019). Based on the *UhAvr1* transcription profiles, the early stages of infection are the likely time points when this effector performs its virulence function and is recognized in cultivars having *Ruh1* (avirulence function). Detailed microscopy at the initial stages of infection upon inoculation of cv. Odessa with UhAvr1m teliospores displayed hyphal death at 48 hpi and a subsequent lower fungal biomass at 72 hpi (Figure 1 and 4B), supporting a role for this effector in suppressing plant defences at these stages. After 72 hpi, hyphal growth from this inoculum seemed to have recovered (Figure 1 and 4B) possibly due to a redundant function from other effector(s) that are working in parallel or consecutively to target the same host protein(s) or same pathway(s), overwhelming the plant defences. For example, two unrelated bacterial effectors from *P. syringae*, AvrRps4 and HopA1, target the central regulator of basal resistance, EDS1 (ENHANCED DISEASE SUSCEPTIBILITY1) [60]. It is hypothesized that the functional redundancy of effectors highlights the importance of a host target for successful colonization [9]. Indeed, individual deletion of *UhAvr1* orthologous gene family members in *Um* (*tin1-1* to *tin1-5*) or deletion of a number of other effector clusters did not reduce virulence on maize [51,61]. Similarly, deletion of *UhAvr1* and its paralogous gene (UHOR_10021) did not compromise virulence of *Uh* during a compatible interaction as measured in the number of diseased plants at 2 to 3 months [16], suggesting the involvement of additional effectors. Although the *UhAvr1* paralogous gene has not been functionally characterized, recent *Uh*-barley leaf transcriptomic data revealed up-regulation of UHOR_10021 from 3 dpi [18]. This time point (72 hpi) coincides with our observed recovery of fungal growth emanating from UhAvr1m teliospores. Further experiments will be needed to confirm a functional redundancy of this paralogue or other effectors playing a role similar to UhAVR1.

Functions of effectors range from shielding hyphae to avoid host recognition, suppression of host defence responses, manipulation of the host physiology for growth and reproduction, and inducing plant cell death in the case of necrotrophic fungi [9,11]. Orthologous gene family members of *UhAvr1* in *Um* (*tin1-1* to *tin1-5*) were implicated in basal defence responses based on a transcriptomic analysis in maize [61]. Similarly, our results suggest at least a function in virulence for UhAVR1 in suppressing (a) defence component(s) involved in basal immunity (Figures 1, 4 and S3). Suppression of nonhost reactions in barley by UhAVR1 delivered via the *Psa* T3SS (Figure 4F-G) suggests that its target(s) may be involved in PTI responses. Also, its ability to suppress three cell death inducers involved in PTI and ETI in the nonhost *Nb* (Figure 4H) suggests that UhAVR1 targets common component(s) of PTI and ETI. Delivery of UhAVR1 in barley using the FoMV effector expression system showed a distinct HR reaction in two *Ruh1* carrying cultivars, Hannchen and SM89010 (Figure 4E) validating the effectiveness of this system in expressing this effector. While testing this system, barley cv. Odessa and heterologous *Nb* showed heightened susceptibility to FoMV upon expression of VOX:UhAvr1+SP (Figures 4C-D and S3). Overall, this suggests the conservation of (a) target(s) for UhAVR1 in both *Nb* and barley. Indeed, several effectors from bacteria, fungi and oomycetes have been shown to suppress immunity-associated cell death when expressed transiently in plant cells [62–66]. For example, effector Shr7 from the wheat stripe rust fungus has been shown to suppress PTI responses triggered by fgl22 in *Nb* and non-specific HR triggered by *P. syringae* DC3000 in wheat plants [65].

Hyphal death and reduced fungal biomass (Figures 1 and 4A) from 48 hpi onwards in cv. Hannchen supports the notion of recognition of this effector or its action by *Ruh1*. Using electron microscopy, our lab had previously demonstrated an effective and very localized defence response in cv. Hannchen elicited by *Uh* isolates having *UhAvr1* at 48 hpi involving the formation of host cell wall appositions at the site of penetration along with degraded mycelia at the site of invasion [2]. The lack of visible macroscopic symptoms of cell death distinctive of an HR in the cv. Hannchen is in congruence with the lack of observed PI stained barley cells, but dead hyphae on the surface during our confocal microscopy observations (Figure 1). Examples of plant defences at the site of fungal penetration leading to none or limited fungal growth exist in other cereal pathosystems [67–70]. For seed-borne smut fungi, an elicited HR response along with fungal arrest has been suggested to be a defence mechanism to delay the fungi in reaching the (barley) crown tissue before floral differentiation occurs [4].

### 4.2. Secretion, translocation and in planta localization of UhAVR1

Successful plant colonization by microbes involves the secretion of effector molecules by a variety of structures. Gram-negative bacteria deliver effectors inside a host cell’s cytoplasm using the well-characterized type III secretion system (T3SS) [10]. Pathogenic fungi and oomycetes are thought to secrete effectors to the apoplast or cytoplasm of its host cells, either using feeding structures such as haustoria or infection structures such as appressoria [11,12], though secretion from hyphae cannot be ruled out. Interestingly, microscopy of barley seedlings infected with *Uh* teliospores revealed a knoblike appressorium structure which was observed in all interactions but not at all infection sites (Figures 2 and S2). Studies performed in barley coleoptiles inoculated with *Blumeria graminis* showed that excessive humidity along with cuticle damage of coleoptiles can affect the maturation and penetration of fungal appressoria [71]. Germination of barley seedlings in Petri dishes at higher humidity and teliospore inoculation of barley coleoptiles using cotton buds in our experiments may have contributed to infection without the formation of visible knob-like structures.

The presence of an N-terminal SP for extracellular secretion is one of the hallmarks of effectors from eukaryotic pathogens [72]. The SP directs a protein to the ER-Golgi where the SP is cleaved off during translocation and the mature protein is packed into vesicles for extracellular export [73]. However, examples of effectors that lack a SP and are secreted via an unconventional pathway exist [74]. In *Magnaporthe oryzae* and *Phytophthora infestans*, a conventional pathway for secretion of apoplastic effectors has been described alongside an unconventional pathway for secretion of effectors that are found in the cytosol [55,75,76]. In our study, the deletion of the *UhAvr1* SP completely blocked the secretion of this effector from the fungus into liquid medium (Figure 5D) supporting the role of this SP in secretion. The sensitivity to BFA of secretion of *UhAvr1* and *UmPit2* from fungal cells supports secretion via the ER-Golgi pathway for both effectors (Figure 6). The cleavage of SP from the UhAVR1 mature protein was supported by the presence of a same molecular weight mature protein produced from UhAVR1-SP:GFP and UhAVR1+SP:GFP in protein blots from agroinfiltrated *Nb* (Figure S6C). This confirms that the SP is cleaved off presumably upon processing and secretion through the ER-Golgi. Indeed, our results show that the *UhAvr1* SP was also able to direct the efficient translocation of another protein (GFP) through the *Nb* secretory pathway as evidenced by the detection of cleaved forms of a GFP on protein blots and protein aggregation under confocal microscopy during BFA assays (Figures S8C and S10).

We showed that the UhAVR1 effector is targeting some conserved components of basal immunity and is secreted via the ER-Golgi pathway but the exact location of this effector in the host is unknown. Apoplastic effectors target surface receptors or extracellular components of resistance [12]. They have also been implicated in protecting fungi from degradation or recognition by the plant immune system [9]. Whereas cytosolic effectors are translocated inside the host cell where they interact with components of cytoplasmic immunity some of which that are coded by *R* genes [9,11,12]. The combination of transient assays in barley and *Nb* resulted in a cytosolic localization for UhAVR1 irrespective of the presence of its SP (Figures 7 to 9 and S5). Similarly in *Nb*, UhAVR1 with or without its SP and tagged either with mCherry or GFP showed a cytosolic localization that does not co-localize with any organelle (Figures 8, 9 and S7). A nucleoplasm localization was seen for UhAvr1-SP:GFP produced under a strong constitutive promoter in *Nb* (Figure 8A). In protein blots, breakdown products were also detected at low levels by anti-GFP antibody (Figure 8C and S10B). The GFP moiety can cross the nuclear pore complex size threshold of 40 kDa [77] so the breakdown product detected by anti-GFP could have entered the nucleus resulting in similar GFP fluorescence observed in control GFP plants. Altogether, these results support the cytosol as the site of the virulence and avirulence function of this effector.

Here, it should be noted that the effectors were produced inside the host cell from T-DNA delivered by agroinfiltration or from the cytoplasmic replicating VOX vectors. Similarly, in the *Pseudomonas* delivery systems, the effectors are first produced in the bacterial cells and then injected into the host cell cytoplasm. These artificial systems place the UhAVR1 (with or without the SP) inside the host cell instead of its likely secretion into the host apoplast by the fungus. In this case, the delivery of UhAVR1 with SP by these artificial systems should allow UhAVR1 to enter the host secretory pathway and exit into the apoplastic space using the host secretory pathway. In our study, no apoplastic localization was seen for UhAvr1+SP:mCherry in *Nb* (Figure 8B). There are two possible scenarios for this: UhAVR1+SP:mCherry does not exit the host cell or it is exiting and re-entering the host cell. In another pathosystem, in addition to secretion from the host cell of mature *P. infestans* avirulence effector AVR3A^KI^, miss-targeting from the ER to the cytosol was suggested to occur due to overexpression in *Nb* [63]. However, we support the notion that UhAVR1+SP gets secreted and reenters the host cell. First of all, translocation of misfolded proteins occurs in the ER [78] which is upstream to the site of action of BFA. In eukaryotes, BFA inhibits the formation of COPI-coated vesicles required for protein passage to the Golgi complex [41] and we have seen protein accumulation of UhAVR1+SP:mCherry or other proteins with SP upon exposure to BFA (Figures 9 and S10) suggesting the effector reaches the Golgi apparatus. Thus, miss-targeting of UhAVR1+SP:mCherry from the ER can be ruled out.

Secondly, we showed that *Ustilago* fungal effectors with a SP are capable of utilizing the plant secretory pathway as evidenced by the secretion of the effector UmPit2+SP:mCherry to the apoplast of *Nb* (Figures 9 and S9). Protein blots showed an increased amount of breakdown products indicating protein degradation, upon exposure of agroinfiltrated UmPit2+SP:mCherry or UhAvr1+SP:mCherry to water and a lesser amount when exposed to BFA, supporting the notion that secretion to the apoplast, a hostile environment with many proteases, could be responsible for this (Figure 9B). Thus, secretion to the apoplast can be inferred to occur for UhAvr1+SP:mCherry where the SP gets cleaved before its secretion. If this holds true, mCherry fluorescence is expected to be seen in the apoplast. However, confocal microscopy is a snapshot taken at the specific time-point. Apoplastic accumulation of the protein at any time point is dictated by the rate of secretion (R_s_) and the rate of re-entry (R_r_) considering other factors remain constant. If the rate of secretion is higher than the rate of re-entry (Rs>Rr), proteins accumulate in the apoplast causing higher apoplastic fluorescence and the intensity difference between cytosol and apoplast would depend on the difference between R_s_ and R_r_. In contrast, if the rate of secretion is equal to the rate of re-entry (R_s_=R_r_), fluorescence will mainly be observed in the cytosol as the protein only transits the apoplast.

Thirdly, during our assays, suppression of nonhost reactions (Figure 4F-G) as well as suppression of cell death inducers (Figure 4H) occurred upon expression of UhAVR1-SP in the cytosol supporting an intracellular site of action. The nonhost reaction in barley was also suppressed upon the delivery of UhAVR1+SP by the *Psa* T3SS (Figure 4F-G) suggesting possible secretion followed by the re-entry of UhAVR1 in the host cytosol to suppress PTI. Very recently, several effectors from *Arabidopsis*-infecting *Fusarium oxysporum* have been shown to target PTI signalling upon re-entry into *Nb* cells [79]. Similarly, the delivery of UhAVR1+SP using the FoMV VOX vector produced necrosis in barley and in *Nb* (Figure 4C-E). FoMV VOX-mediated delivery of necrotrophic effector ToxA+SP from *Parastagonospora nodorum* into ToxA-sensitive wheat (having toxin sensitivity gene *Tsn1*) led to necrosis [21]. Previous work showed that ToxA is imported into the cells of sensitive wheat in the absence of the pathogen where it localizes to the cytosol and chloroplasts [80]. Similarly, the production of UhAVR1 by VOX:UhAVR1+SP followed by secretion and internalization back into host cells in the absence of the pathogen can be expected. Pathogen-independent re-entry has also been shown for several cytoplasmic rust fungus effectors [81–83], though in some other pathosystems, reentry of effectors may need the presence of the fungus or other fungal factors [84]. However, the lack of visible symptoms in barley and *Nb* upon expression from the VOX:UhAVR1-SP construct (Figure 4C-E) cannot be explained at this time. It is possible that the response of VOX-delivered UhAvr1-SP was subtle and delayed in barley as seen for VOX-delivered effector ToxA-SP [21]. The same holds for *Nb*, if the symptoms were delayed for VOX-delivered UhAVR1-SP then the effects will manifest themselves slowly and may get confounded by the pathogenic effects caused by the viral vector (Figure S3).

## 5. Conclusions

In summary, here we have elucidated the virulence role of UhAVR1 in suppressing PTI and ETI during the early stage of fungal infection and have further characterized the triggering of defence responses in the resistant interaction. This effector is secreted from the fungus via the ER-Golgi pathway and localizes to the host plant cytosol but lacks any previously defined conserved motifs found in other fungi and oomycetes implicated in translocation into plant cells. However, an RFIYL motif in UhAVR1 starting at position 93 was found. The R and L amino acids are conserved in the orthologues in the various smut fungi while the amino acids FIY are interchangeable with other hydrophobic amino acids. Hydrophobic patches of amino acids present in the AvrM effector of the flax rust fungus are required for translocation into the host cell [85]. Further studies will be required to identify the region or the amino acid sequences of the protein involved in the interaction with host proteins resulting in suppressing or triggering defence responses and uptake into the host cell.

## Supplementary Materials

**Figure S1.** Constructs used during this study.

**Figure S2.** Fungal structure observed during the infection with *Ustilago hordei* teliospores on barley coleoptiles.

**Figure S3.** Symptoms on the leaves of *Nb* and barley plants expressing VOX constructs at later stages. **Figure S4.** GFP fluorescence from *Uh* strains expressing GFP tagged UhAVR1 localize to the intercellular space of cv. Odessa cells.

**Figure S5**. UhAVR1:GFP localizes to the cytosol of cv. Odessa cells at 48 hpi.

F**igure S6.** Transcript and protein accumulation of UhAVR1 is affected by the presence of a SP and the SP is cleaved off from the pre-protein in *Nb*.

**Figure S7.** UhAVR1 does not co-localize with plant organelle markers.

**Figure S8**. GFP expressed with the N-terminal SP of UhAVR1 localizes to the cytosol of *Nb* at 24 hpi. **Figure S9**. UmPit2:mCherry localizes to the apoplast in leaves of *Nb*.

**Figure S10**. Brefeldin A exposure in *Nb* blocks trafficking of proteins with the UhAVR1 SP. **Table S1.** Fungal strains used in this study.

**Table S2.** Oligonucleotides used in this work.

**Table S3.** *UmPit2* effector mRNA sequence with SP (underlined) and STOP codon.

**Table S4.** Peptide sequence used to generate the UhAVR1 antibody.

**Table S5.** Pathogenicity assays of N-terminally GFP tagged *UhAvr1* strains showed loss of avirulence.

## Author Contributions

Conceptualization, A.P.M.A. and G.B.; experimentation and formal analyses, A.P.M.A., S.A., X.S. R.L. and G.B.; funding acquisition, G.B.; writing—original draft, A.P.M.A.; writing—review and editing, S.A., X.S. and G.B. All authors have read and agreed to the published version of the manuscript.

## Funding

This research was funded in part by the Natural Sciences and Engineering Research Council of Canada, grant number RGPIN-2015-05738.

## Acknowledgments

We are grateful to Peter Moffett (U. of Sherbrooke, Canada), Diane Cuppels (AAFC, ON, Canada), Brian J. Staskawicz (U. of California, USA), Kostya Kanyuka (Rothamsted Research, UK) and Sophien Kamoun (The Sainsbury lab, UK) for providing P-bodies markers, the *Psa* strain, vector pPSV_PsSPdes, the FoMV vectors and INF1 expression construct, respectively. We thank Pankai Bhowmik for an improved protocol and advice on generating barley protoplasts.

## Conflicts of Interest

The authors declare that they have no conflict of interest. The funders had no role in the design of the study; in the collection, analyses, or interpretation of data; in the writing of the manuscript, or in the decision to publish the results.

## Notes

### Competing Interest Statement

The authors have declared no competing interest.

