## Supplementary Materials for "UhAVR1, an HR-triggering avirulence effector of *Ustilago hordei*, is secreted via the ER-Golgi pathway to the cytosol of barley coleoptile cells and contributes to virulence early in infection"

### Supplementary Figures

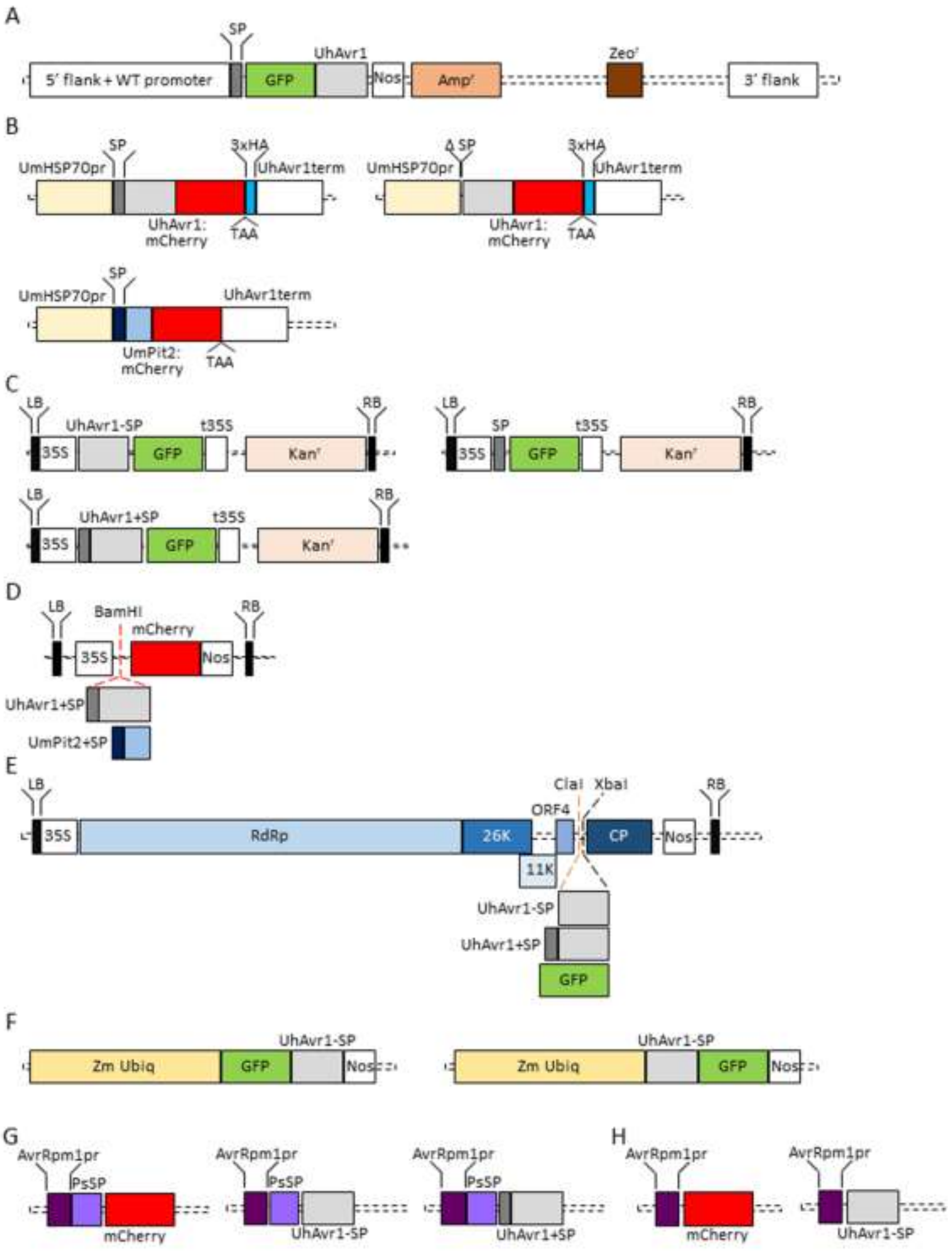


**Figure S1. Constructs used during this study. (A)** Gene cassette SP:GFP:UhAvr1:Nos used to generate the strain Uh1398. A gene cassette SP:GFP:UhAvr1:Nos was cloned downstream of sequences for homologous recombination (5′

flank + WT promoter and 3′ flank). **(B)** FunGus expression vectors used in this study expressing UhAvr1+SP:mCherry, ΔSP:UhAvr1-SP:mCherry or UmPit2+SP:mCherry. **(C)** GFP constructs used in barley and *Nb* transient assays expressing UhAvr1-SP:GFP, UhAvr1+SP:GFP or SP:GFP. **(D)** mCherry constructs used in *Nb* transient assays expressing mCherry, UhAvr1+SP:mCherry or UmPit2+SP:mCherry. **(E)** FoMV-based VOX system showing the organization of open reading frames (ORFs) for viral proteins and target genes UhAvr1-SP, UhAvr1+SP or GFP. **(F)** Constructs used for cell death assays in *Nb* expressing GFP:UhAvr1-SP or UhAvr1-SP:GFP. **(G)** Constructs used for *Psa*-barley assays expressing mCherry, UhAvr1-SP or UhAvr1+SP. **(H)** Deleted *Pseudomonas* SP (PsSP) constructs used for *Psa*-barley assays expressing ΔPsSP:mCherry or ΔPsSP:UhAvr1-SP. Abbreviations: Ampicillin resistance gene ($Amp^r$), Zeocin resistance gene ($Zeo^r$), nopaline synthase terminator (Nos), *Ustilago maydis* constitutive HSP70 promoter (UmHSP70pr), UhAvr1 terminator (UhAvr1term), 3x HA tag (3xHA), stop codon (TAA), left border (LB), right border (RB) for *Agrobacterium* T-DNA transfer, Kanamycin resistance gene ($Kan^r$), CaMV 35S promoter (35S), CaMV 35S terminator (t35S), viral polymerase (RdRp), viral movement proteins (26K, 11K and ORF4), viral coat protein (CP), maize ubiquitin promoter (Zm Ubiq) and *Pseudomonas AvrRpm1* promoter (AvrRpm1pr).

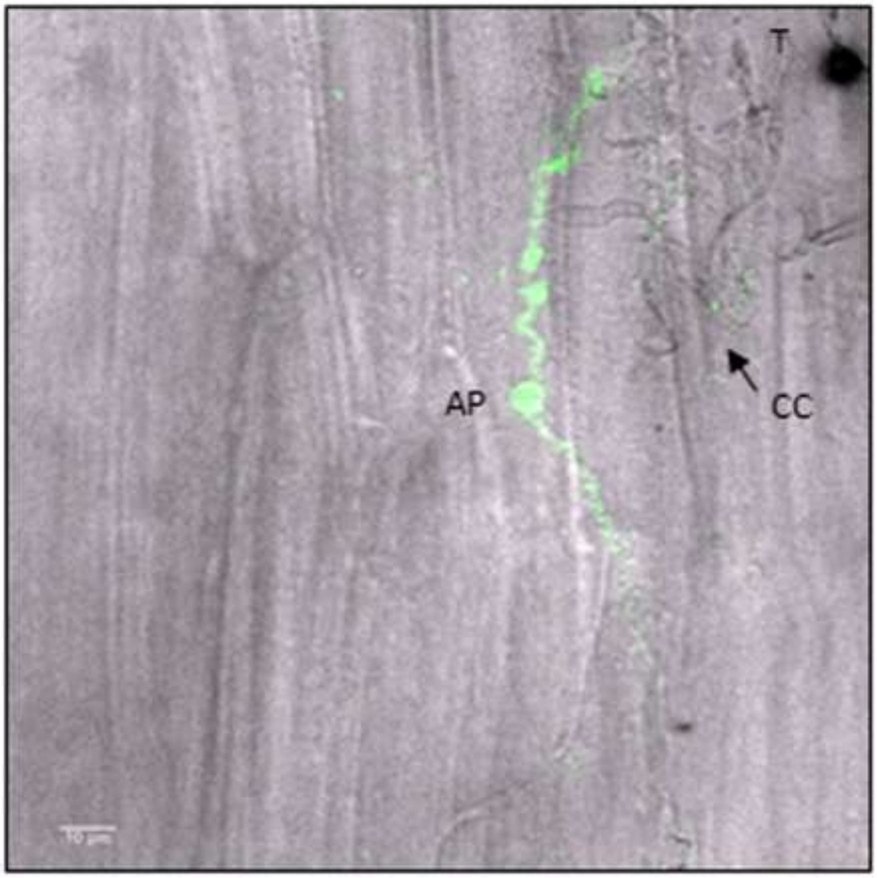


**Figure S2. Fungal structure observed during the infection with *Ustilago hordei* teliospores on barley coleoptiles.** Barley coleoptiles were infected with UhAvr1m teliospores, stained with WGA-AF488 and PI and visualized with confocal microscopy. An appressorium-like (AP) structure was seen near the junction between two neighbouring barley cells. Fungal hyphae with collapsed cells (CC) and a black teliospore (T) were also visible near the appressorium-like structure. Image is a snapshot of a single z-stack observed at 96 hpi in cv. Hannchen and scale bar represents 10 μm.

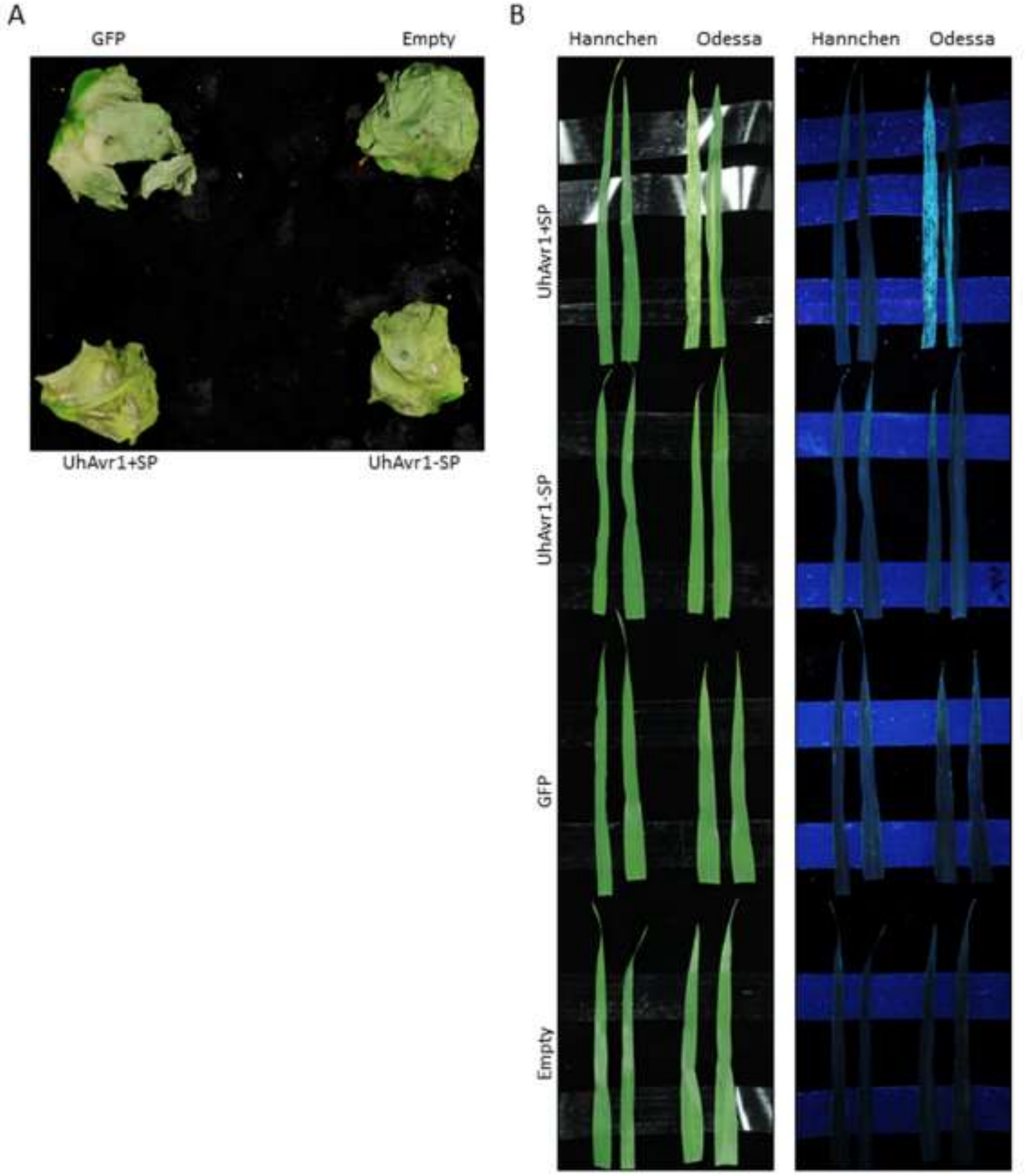


**Figure S3. Symptoms on the leaves of *Nb* and barley plants expressing VOX constructs at late stages. (A)** Agro-infiltrated leaves of *Nb* with VOX constructs expressing GFP, UhAvr1-SP, UhAvr1+SP or an empty-vector control showed necrosis at 13 dpi. The experiment was repeated three times with similar results and a representative image is shown. **(B)** Symptom development on the L3 leaves of barley. L1 and L2 leaves of barley cultivars were rub-inoculated with *Nb* leaf sap previously infected with the VOX constructs as indicated. At 15 dpi, a few plants of cv. Odessa expressing UhAvr1+SP showed mosaic symptoms on their L3 leaves. The images were taken under white light (left) or long-wavelength UV light (right) from the same set of leaves. The experiment was repeated three times with similar results and a representative image is shown.

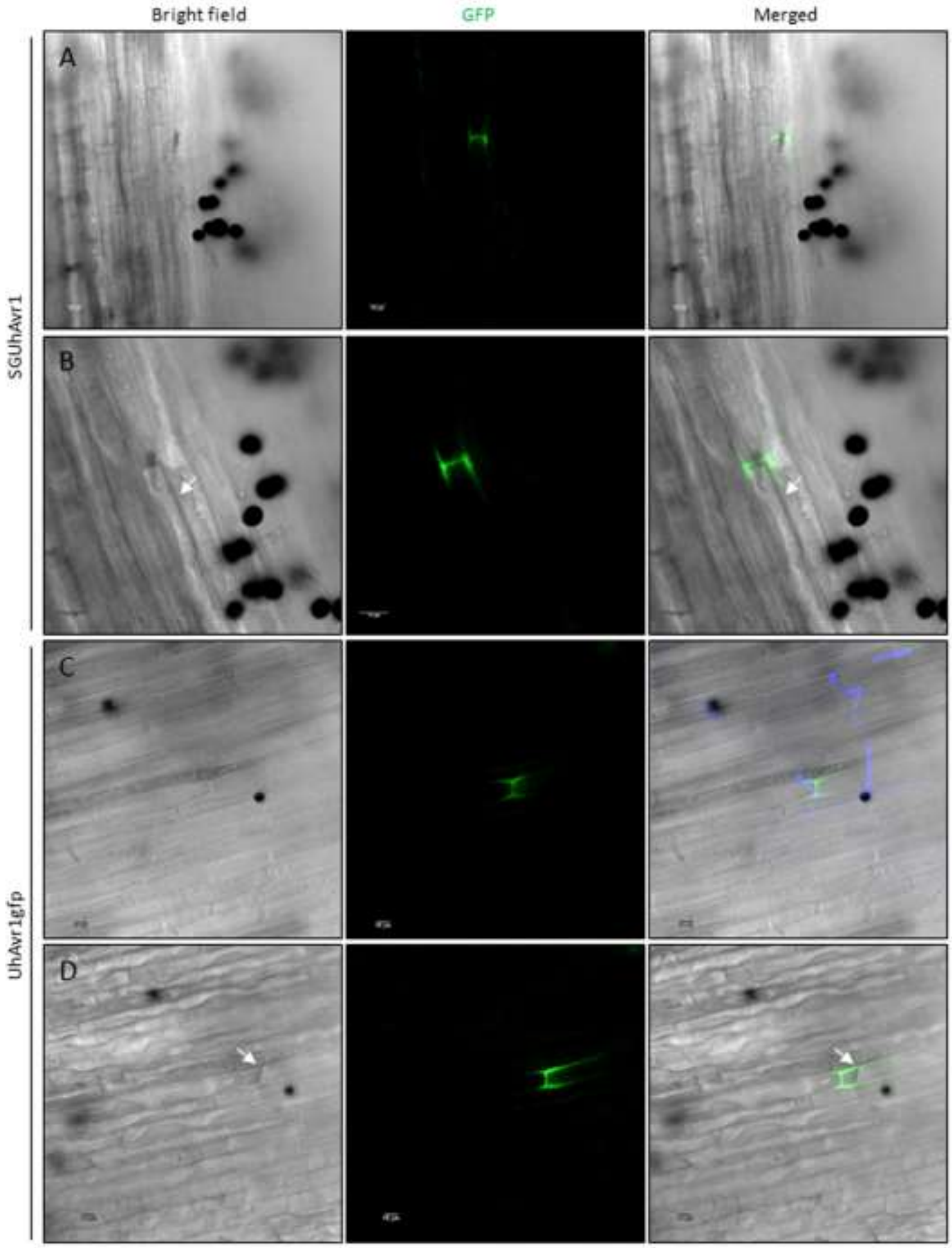


**Figure S4. GFP fluorescence resulting from inoculation with *Uh* strains expressing GFP-tagged UhAVR1 localizes to the intercellular space of cv. Odessa cells.** Confocal imaging of cv. Odessa seedlings infected with SGUhAvr1 **(A and B)** or UhAvr1GFP **(C and D)** teliospores expressing N-terminally or C-terminally GFP-tagged UhAVR1, respectively, revealed the GFP fluorescence in the intercellular space of barley cells near the infection sites. Fungal hyphae were stained with Uvitex 2B (purple) in panel C. **(B and D)** GFP fluorescence remained associated with the cell border even after performing plasmolysis with 1-1.5M NaCl (arrow points to the retracted plasma membrane). All images are snapshots of a single optical section and the scale bar represents 10 μm. Two experimental repeats were performed for both teliospores infections with similar localization results.

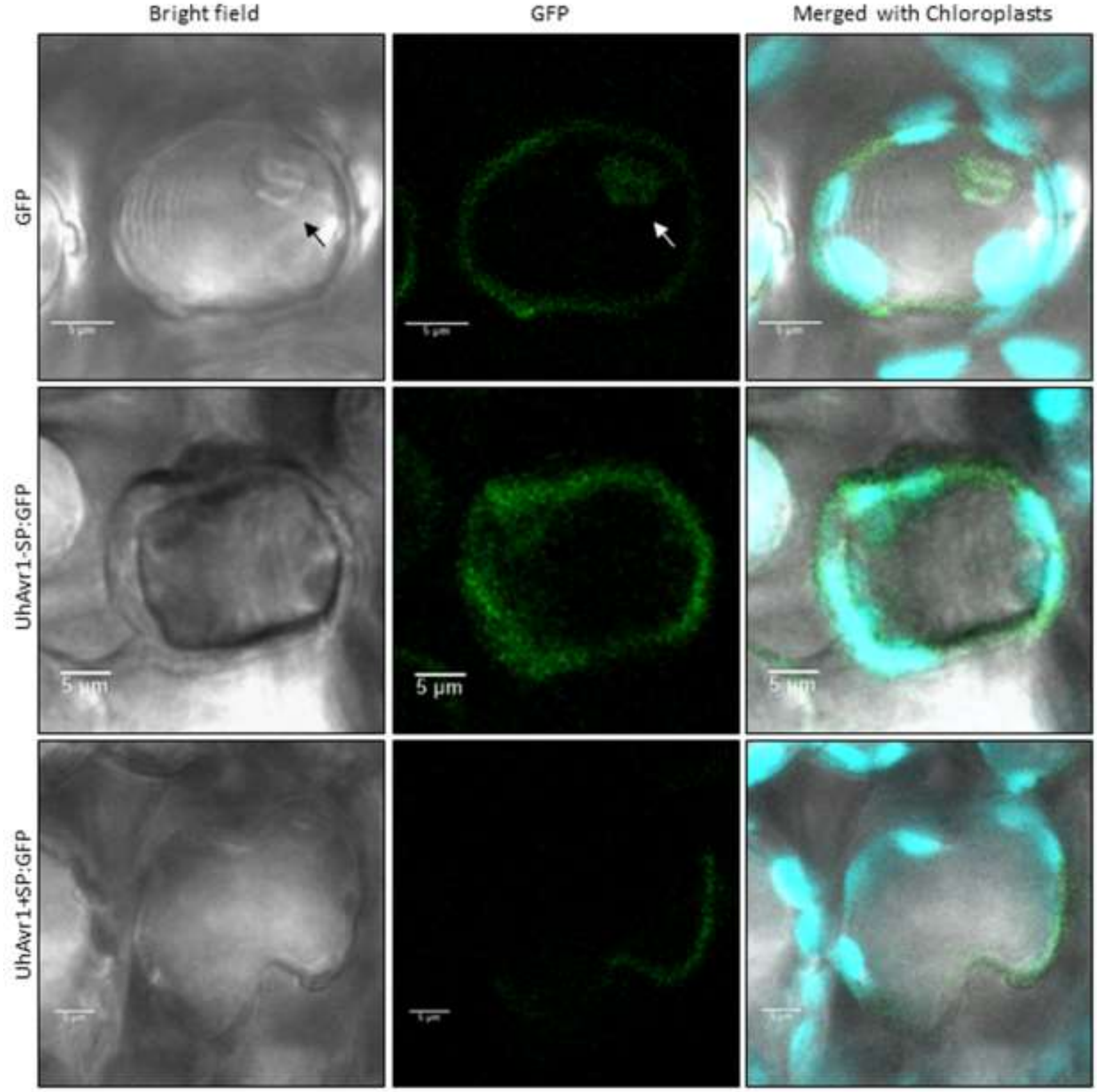


**Figure S5**. **UhAVR1:GFP localizes to the cytosol of cv. Odessa cells at 48 hpi.** Confocal microscopy of leaves of cv. Odessa agroinfiltrated with UhAvr1-SP:GFP or UhAvr1+SP:GFP displayed GFP fluorescence localizing to the cytosol, surrounding the chloroplasts. Whereas, the free GFP control showed fluorescence localizing to the nucleoplasm (arrow) and the cytosol. The same GFP constructs were used for barley protoplast transfection (Figure 7) and *Nb* agro-infiltrations (Figure 8). All images are snapshots of a single optical section and the scale bar represents 5 µm. Confocal imaging of cv. Odessa was performed independently two times since low expression made confocal imaging challenging.

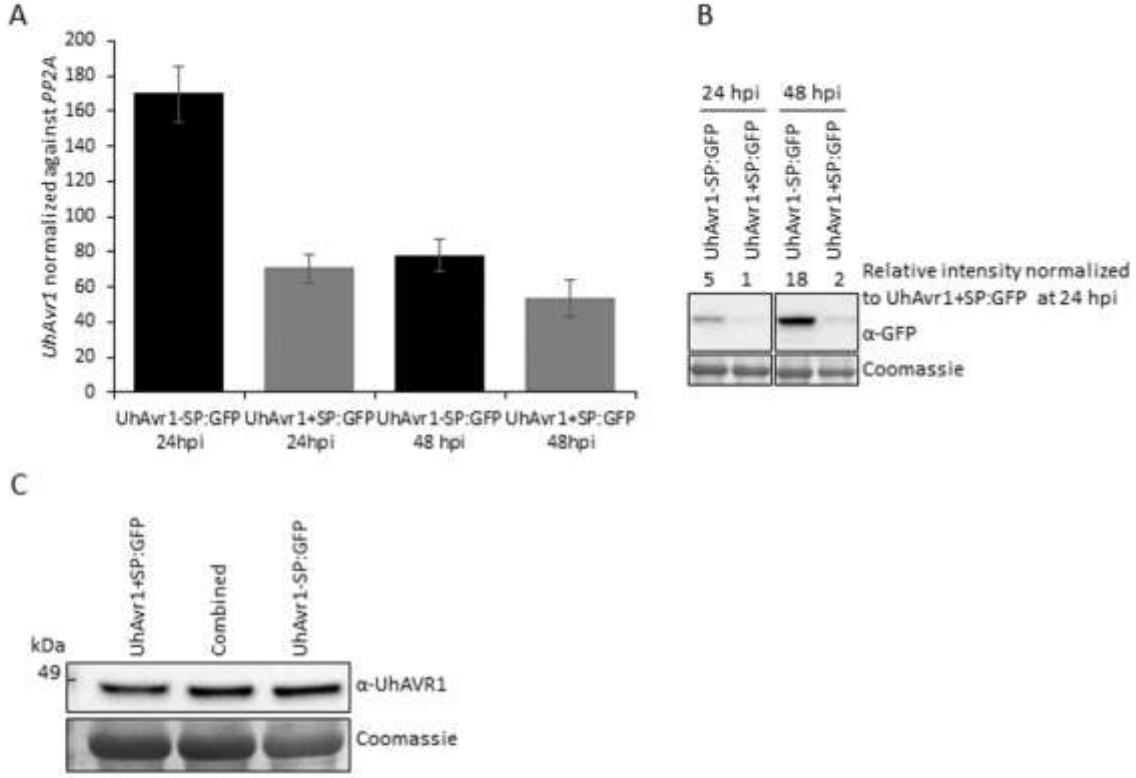


**Figure S6. Transcript and protein accumulation of UhAVR1 is affected by the presence of a SP and the SP is cleaved off from the pre-protein in *Nb*. (A)** Quantification of *UhAvr1* transcripts by ddPCR from agroinfiltrated *Nb* plants expressing UhAvr1-SP:GFP and UhAvr1+SP:GFP at 24 and 48 hpi, with corresponding protein accumulation from one of the experimental repeats (B). Normalization of *UhAvr1* transcripts was done against the *Nb* gene *PP2A*. The ddPCR graph shows the average of two experimental repeats, each including three technical repeats, along with their standard deviation depicted as error bars. **(B)** Protein blot analysis on agro-infiltrated *Nb* at 24 and 48 hpi, corresponding to one of the repeats used in panel A. The intensity value of each protein band detected by anti-GFP antibody was calculated by Image Lab software, normalized to UhAVR1+SP:GFP expression at 24 hpi and is provided on the top of the gel picture. Lower levels of protein as well as transcripts from UhAvr1+SP:GFP compared to UhAvr1-SP:GFP were detected at 24 and 48 hpi. **(C)** Products resulting from the agroinfiltration of UhAvr1+SP:GFP and UhAvr1-SP:GFP were loaded individually (left and right lane) or combined (middle lane) to determine the possible size differences between these two proteins. The sample volumes were adjusted in all three lanes by adding protein lysates from a healthy *Nb* plant to avoid discrepancies in migration of proteins due to unequal protein amounts loaded. No double bands were detected in the combined lane suggesting the presence of the same size protein product, cleaved, mature UhAVR1-SP:GFP.

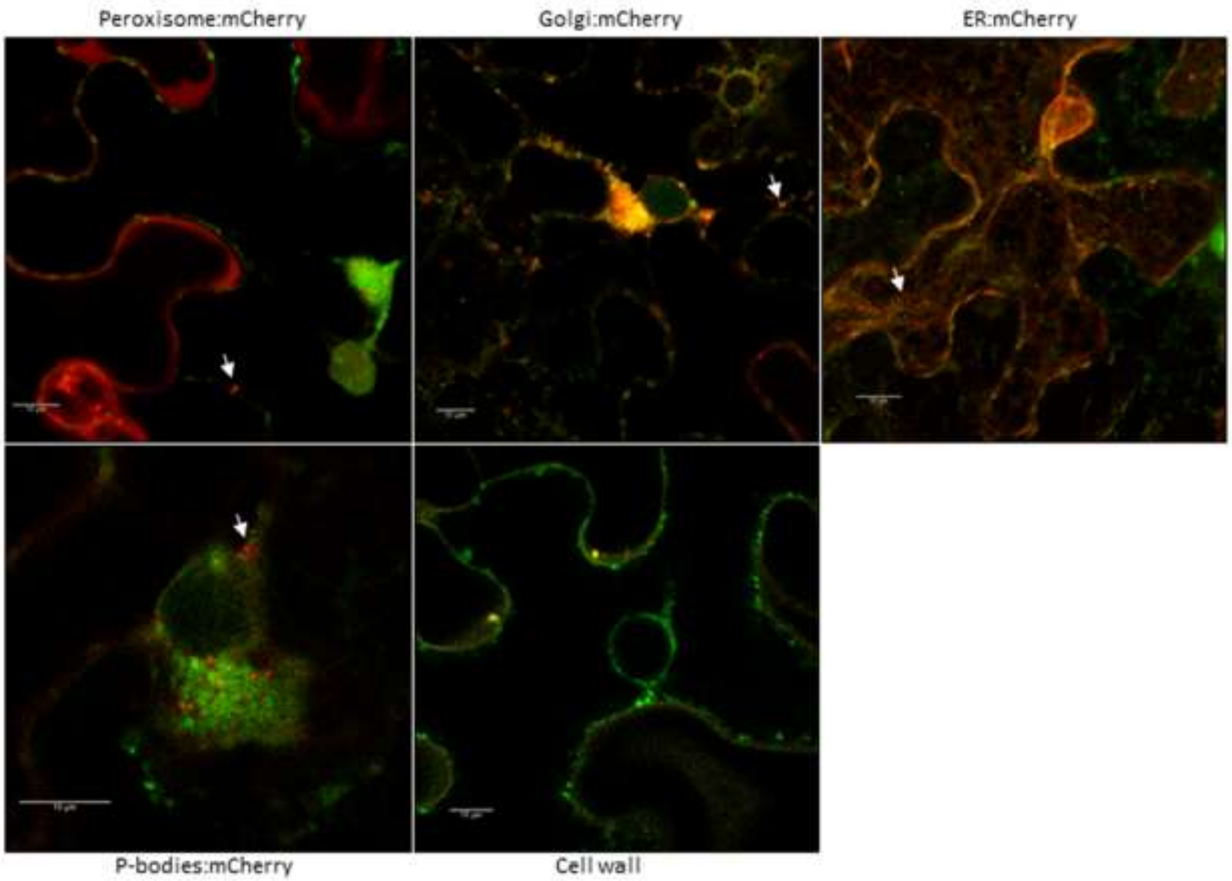


**Figure S7. UhAVR1 does not co-localize with plant organelle markers.** mCherry-tagged organelle markers (arrows) were co-expressed with UhAvr1+SP:GFP via agroinfiltration in *Nb* followed by confocal microscopy. UhAvr1+SP:GFP did not co-localize with the following markers at 24 hpi: peroxisome, golgi, endoplasmatic reticulum (ER) or P-bodies. Visualization of the plant cell wall (yellow) was done at 48 hpi by staining tissues with 0.8 mg/ml PI for 15 min. UhAvr1+SP:GFP fluorescence is seen inside the cell wall. Two independent experimental repeats were performed with the mentioned markers. All images are snapshots of a single optical section except for the ER image which is a maximum projection of a z-stack. Scale bar represents 10 μm.

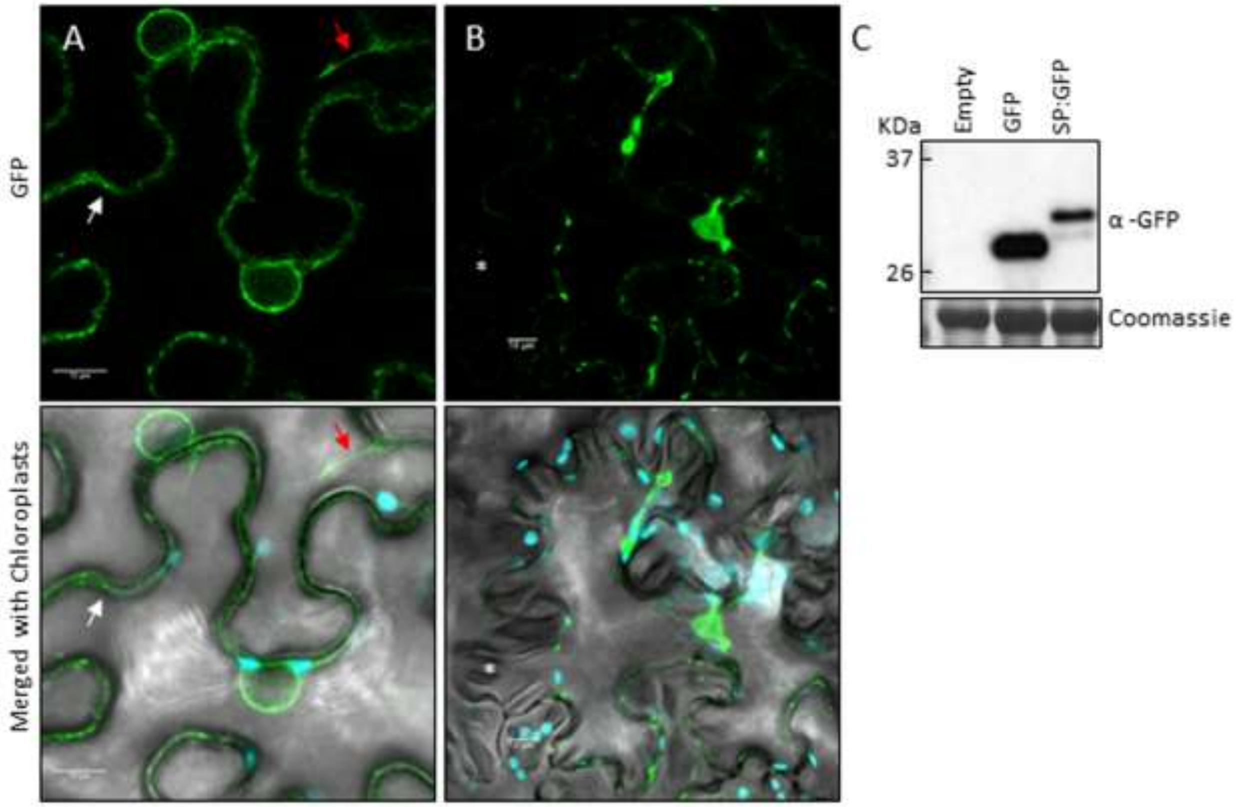


**Figure S8**. **GFP expressed with the N-terminal SP of UhAVR1 localizes to the cytosol of *Nb* at 24 hpi. (A)** Confocal microscopy of *Nb* leaves agroinfiltrated with SP:GFP showed GFP fluorescence localizing to the cytosol (red arrow) and cytosolic foci of different sizes (white arrow). **(B)** No fluorescence was seen in the apoplastic space of *Nb* (asterisk) after plasmolysis was induced with 1.5M NaCl for 20 min. **(C)** Protein blot analysis performed using proteins from the same batch of plants used for panel A showed expressed GFP. Both full length (31 kDa) and cleaved (29 kDa) products were detected from the agro-infiltrated SP:GFP construct. A minimum of three independent repeats were performed and representative results are shown. All images are snapshots of a single optical section and the scale bar represents 10 μm.

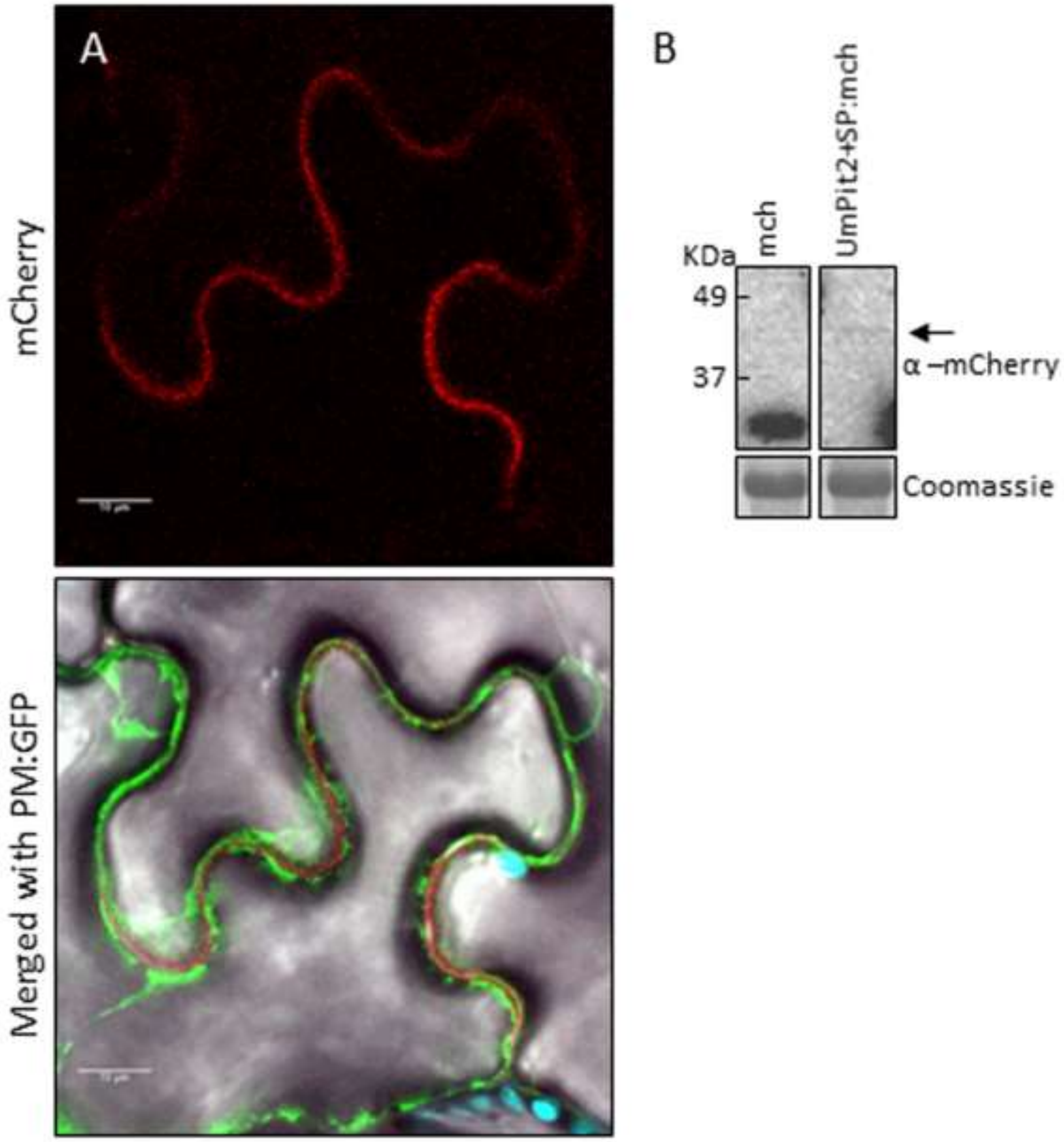


**Figure S9**. **UmPit2:mCherry localizes to the apoplast in leaves of *Nb*.** Leaves of *Nb* plants were co-agroinfiltrated with constructs expressing UmPit2+SP:mCherry and a GFP tagged plasma membrane (PM) marker. **(A)** Confocal microscopy at 24 hpi showed the mCherry fluorescence localizing in between the GFP signals. Images are a snapshot of a single optical section and scale bar represents 10 µm. **(B)** Protein blot analysis of the samples isolated from the same batch of plants used in panel A showed the presence of mCherry and SP-cleaved UmPit2:mCherry (arrow). The image originates from the same blot. A representative result from a minimum of three independent repeats is shown.

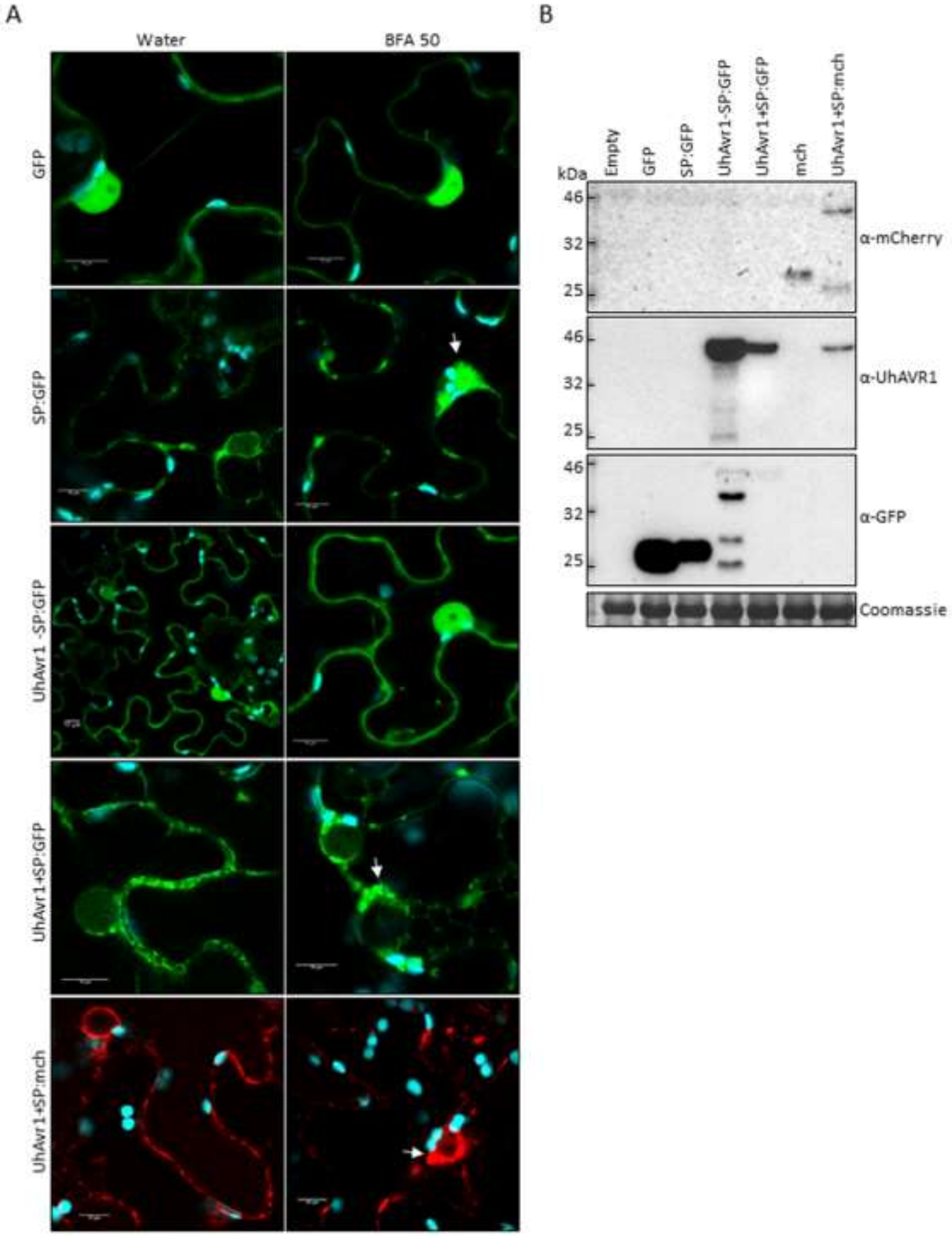


**Figure S10**. **Brefeldin A exposure in *Nb* blocks trafficking of proteins with the UhAVR1 SP. (A)** *Nb* plants transiently expressing fluorescence tagged proteins were exposed to BFA or water. Confocal imaging showed that the constructs with a signal peptide (SP:GFP, UhAvr1+SP:GFP and UhAvr1+SP:mCherry) presented protein aggregation (arrows) only upon exposure to BFA. Constructs lacking a SP (GFP and UhAvr1-SP:GFP) did not show protein aggregation upon water or BFA exposure. All images are snapshots of a single optical section and the scale bar represents 10 μm. **(B)** Protein blot analysis of proteins isolated from the plants used in panel A, prior to Brefeldin A or water exposure, showed the presence of mainly full-length proteins. A representative result from two independent repeats is shown for confocal and protein blots.

**Table S1. Fungal strains used in this study**

| Strain ID | Genotype | Comments |
|---|---|---|
| Um001 | *a2b2* | Wild type *U. maydis* strain, alias Um518; [1]. |
| Um002 | *a1b1* | Wild type *U. maydis* strain, alias Um521; [1]. |
| Uh362 | *MAT-2 Uhavr1* | Wild type *U. hordei* strain, alias Uh4854-10; [2]. |
| Uh364 | *MAT-1 UhAvr1* | Wild type *U. hordei* strain, alias Uh4857-4; [2]. |
| Uh1289 | Uh364 (*MAT-1 ΔUhAvr1*), carboxin$^r$ | *UhAvr1* deletion strain, [3]. |
| Uh1351 | Uh364 (*MAT-1 UhAvr1 [otef:gfp]*), zeocin$^r$ | Control strain that constitutively expresses a genome-integrated otef:GFP; [3]. |
| Uh1353 | Uh1289 (*MAT-1 ΔUhAvr1* [*UhAvr1:GFP*]), zeocin$^r$ | *UhAvr1* deleted strain expressing *UhAvr1:GFP* from the *UhAvr1* promoter. Chimer is located in *UhAvr1* endogenous location; [3]. |
| Uh1357 | Uh1289 (*MAT-1 ΔUhAvr1* [*otef:UhAvr1:GFP*]), zeocin$^r$ | *UhAvr1* deleted strain expressing *UhAvr1:GFP* from the otef promoter. Chimer is located in *UhAvr1* endogenous location; [3]. |
| Uh1397 | Uh1289 (*MAT-1 ΔUhAvr1* [*SP:GFP:UhAvr1*]), zeocin$^r$ | *UhAvr1* deleted strain expressing *SP:GFP:UhAvr1* from its *UhAvr1* promoter. Chimer is located in *UhAvr1* endogenous location. Clone 1. This work. |
| Uh1398 | Uh1289 (*MAT-1 ΔUhAvr1* [*SP:GFP:UhAvr1*]), zeocin$^r$ | *UhAvr1* deleted strain expressing *SP:GFP:UhAvr1* from its *UhAvr1* promoter. Chimer is located in *UhAvr1* endogenous location. Clone 2. This work. |
| Uh1399 | *Uh1289 (MAT-1 ΔUhAvr1 [SP:GFP:UhAvr1]), zeocin*$^r$ | *UhAvr1* deleted strain expressing *SP:GFP:UhAvr1* from its *UhAvr1* promoter. Chimer is located in *UhAvr1* endogenous location. Clone 3. This work. |
| Uh1430 | Uh1351 (*MAT-1 UhAvr1 [UhAvr1+SP:mCherry]*), carboxin$^r$ zeocin$^r$ | A strain expressing *UhAvr1+SP:mCherry* in pUHESdest episomal plasmid along with genome-integrated otef:GFP. This work. |
| Uh1434 | Uh1351 (*MAT-1 UhAvr1 [ΔSP:UhAvr1:mCherry]*), carboxin$^r$ zeocin$^r$ | A strain expressing *ΔSP:UhAvr1:mCherry* in pUHESdest episomal plasmid along with genome-integrated otef:GFP. The SP was deleted from pUHESdest vector backbone. This work. |
| Uh1440 | Uh1351 (*MAT-1 UhAvr1 [UmPit2+SP:mCherry]*), carboxin$^r$ zeocin$^r$ | A strain expressing *UmPit2+SP:mCherry* with its SP in pUHESdest episomal plasmid along with genome-integrated otef:GFP. This work. |
| WT teliospores | Uh362 x Uh364 (*MAT-2 MAT-1 Uhavr1 UhAvr1*) | This work. |
| UhAvr1m teliospores | Uh362 x Uh1289 (*MAT-2 MAT-1 Uhavr1 ΔUhAvr1*) | This work. |
| GFP teliospores | Uh362 x Uh1351 (*MAT-2 MAT-1 Uhavr1 UhAvr1 [otef :gfp]*) | Described in [3]. |
| SGUhAvr1 teliospores | Uh362 x Uh1398 (*MAT-2 MAT-1 Uhavr1 ΔUhAvr1* [*SP:GFP:UhAvr1*]) | *SP:GFP:UhAvr1* is driven by UhAvr1 wild type promoter. This work. |
| UhAvr1GFP teliospores | Uh362 x Uh1357 (*MAT-2 MAT-1 Uhavr1 ΔUhAvr1* [*otef:UhAvr1:GFP*]) | *SP:GFP:UhAvr1* is driven by otef constitutive promoter; [3]. |

**Table S2. Oligonucleotides used in this work**

| No. | Primer Name | Sequence (5′-3′) | Purpose |
|---|---|---|---|
| 520 | Ta_ubi-conj_fw | AGCATTTGCCTTGACATTCTCA | Primer pairs for amplification of barley reference gene *HvUbiq* (*ubiquitin-conjugating enzyme-specific*). |
| 521 | Ta_ubi-conj_rev | CCCGATCAGTCTTGTACATGTGA | |
| 1247 | UH_10022_fw | CACCATGCGATCGTTTTCCCTTTTCC | Forward primer to generate entry vector of the SP of *UhAvr*1. In red are CACC sequences for directional cloning into the GateWay™ entry vector. |
| 1248 | UH_10022-SP_Fw | CACCATGCCTGGCGACAAAGCTTCTTC | Forward primer to generate entry clone UhAvr1-SP:mCherry. In red are CACC sequences for directional cloning into the GateWay™ entry vector. |
| 1504 | mCherry_fw | CACCATGGTGAGCAAGGGCGAGGAGG | Primer pairs to generate entry clone mCherry. In red are CACC sequences for directional cloning into the GateWay™ entry vector. |
| 1505 | mCherry_rev | CTTGTACAGCTCGTCCATGCC | |
| 1764 | recombined pVSP EcoRV site 1_fw | CCGGATATCACAAGTTTG | Primers to generate *Pseudomonas* vector pVSPΔSPΔRFC-A, where the *Pseudomonas* SP and GateWay™ cassette A have been deleted. Restriction sites are underlined. |
| 1765 | recombined pVSP EcoRV site1_rev | CCGGATATCACCACTTTGTAC | |
| 1770 | pVSP frag1 5′ BglII_fw | GAAGATCTGTAGATTTGTGATGG | |
| 1771 | pVSP frag2 3′ BglII_rev | GAAGATCTTGATCCCCTG | |
| 1776 | pVSP-EcoRV frag 2 5′EcoRV_fw | GATCGATATCGGATCGATCCTATCCGTACG | |
| 1777 | pVSP-EcoRV frag 1 3′EcoRV_rev | CGCGATATCCATAAAAAACC | |
| 1794 | SalI_3′ flank Uh10022_fw | GGCGTCGACCTTAGCCTAGTCCCGCTCT | 3′ flanking primer to generate *UhAvr1* gene probe. SalI site is underlined |
| 1795 | SpeI_3′ flank Uh10022_rev | GGCACTAGTGAGAAGAAGCAGGGCTTTCA | 3′ flanking primer to generate *UhAvr1* gene probe. SpeI site is underlined |
| 1965 | UhGapdh-fw | CAAGGCTCAGATCGTCTCCA | Primer pairs to amplify *Uh* fungal reference gene *UhGapdh* (*glyceraldehyde 3-phosphate dehydrogenase*) |
| 1966 | UhGapdh-rev | GGATGATGTTGGCAGCAGCG | |
| 1967 | Uh10022_ddPCR1_fw | AGTAGACAGGAATCTTGCCTGG | Primer pairs to quantify *UhAvr1* expression in ddPCR |
| 1968 | Uh10022_ddPCR1_rev | GGTCAGAACGTCTCCAATCTCG | |
| 2031 | BamHI_Avr1+SP_fw | GGAGGCGGATCCAACCATGCGATCGTTTTCCCTT | Primer pairs to clone *UhAvr1+SP* into pCAMBIA:mCherry vector. BamHI site is underlined |
| 2032 | Avr1+SP_BamHI_rev | TAAAACGGATCCACCTCCTCCGGCAAATCGGAGCGC | |
| 2034 | mCherry_STOP_rev | **TTA**CTTGTACAGCTCGTCCATGCC | Reverse primer to generate entry clone UhAvr1-SP:mCherry. The natural stop codon is in bold. |
| 2035 | SPUhAvr1_rev | AGA AGC TTT GTC GCC AGG TGC | Reverse primer to generate entry vector of the SP of *UhAvr1*. |
| 2098 | pUHes_UhAvr1mCherryΔSP_fw | GGCGAATTCTGATATCACAAGTTTGTACAAAGCAGG | |

| | | | |
|---|---|---|---|
| 2099 | pUHes_UhAvr1mCherryΔSP_rev | CCGGAATTCATTGCCCCCCGGGATCTG | Outward-facing primers to delete the SP of the pUHESdest vector backbone by inverse PCR. EcoRI site is underlined. |
| 2125 | pUHES_gib_fw | TGACGCGCGGTTTGACTTGTC | Primer pairs to amplify pUHESdest for cloning *UmPit2+SP:mCherry* via Gibson assembly. |
| 2126 | pUHES_gib_rev | TGCCCCCCGGGATCTGGC | |
| 2127 | UmPit2+SPmCherry_gib_fw | TTGCCAGATCCCGGGGGGCA**ATG**CTGTTTCGCTCAGCC | Primer pairs to amplify and clone *UmPit2+SP:mCherry* into pUHESdest via Gibson assembly. Start and stop codons are in bold. |
| 2128 | UmPit2+SP mCherry_gib_rev | ACAAGTCAAACCGCGCGTCA**TTA**CTTGTACAGCTCGTCCATG | |
| 2129 | BamHI_UmPit2+SP_fw | GGAGGCGGATCCAACC**ATG**CTGTTTCGCTCAGCC | Primer pairs to clone *UmPit2+SP* into pCAMBIA: mCherry vector. BamHI site is underlined and the start codon is in bold. |
| 2130 | UmPit2+SP_ BamHI_rev | TAAAACGGATCCACCTCCTTCCCAGATGACCACATCTCCGT | |
| 2174 | GFP5'-ClaI-fw | AGGTCAATCGAT**ATG**GTGAGCAAGGGCGAGG | Primer pairs to clone GFP into FoMV PV101. ClaI and XbaI site is underlined. Start and stop codon are in bold. |
| 2175 | GFP3'-XbaI-rev | TATGCTTCTAGA**TTA**CTTGTACAGCTCGTCCATG | |
| 2176 | UhAVR1_Cla1-fw | AGGTCAATCGAT**ATG**CGATCGTTTTCCCTTTTCCTC | Forward primer to clone *UhAvr1+SP* into FoMV PV101. ClaI site is underlined and the start codon is in red. |
| 2177 | UhAVR1-SP _Cla1-fw | AGGTCAATCGAT**ATG**CCTGGCGACAAAGCTTC | Forward primer to clone *UhAvr1-SP* into FoMV PV101. ClaI site is underlined and the start codon is in red. |
| 2178 | UhAVR1_XbaI-rev | TATGCTTCTAGA**TCA**TCCGGCAAATCGGAGCG | Reverse primer to clone *UhAvr1* into FoMV PV101. XbaI site is underlined and the natural stop codon is in red. |
| 2224 | Nb_PP2A_fw | GACCCTGATGTTGATGTTCGCT | Primer pairs to amplify *Nicotiana benthamiana* reference gene *PP2A* (*protein phosphatase 2*). |
| 2225 | Nb_PP2A_rev | GAGGGATTTGAAGAGAGATTTC | |

**Table S3.** ***UmPit2*** **effector mRNA sequence with SP (underlined) and STOP codon**
AtgctgtttcgctcagcctttgttctgctcatcgtggcctttgcaagtgcatgcctggtgcaacatgttcaagctattccggtgcgtcgatcgctctctaccgatgcctcaatgagctcggctgctggcaagctcaaccggagatggtggttcggcttcacaggttcgctcggcaaggaacctgacaacggccaagtacagatcaagatcatcccagacgcgctcatcatcaagaatccgcctgccaacaaagacgatctgaacaagctaatcgaaaacctaaaacgcaagcacccaagattcaagacggtggtcatgccgacagatcctaacggagatgtggtcatctgggaaTAA

**Table S4. Peptide sequence used to generate the UhAVR1 antibody**
Anti-UhAVR1 antibody was produced from only 155 amino acids sequences of UhAVR1:
PSFKLEIAENPNVDPFLEKISKLGNSHDLYPHVALMRTTLYGKDKLTTNLGAYPDFRRFIYLGNSPGVPEMYFAVPLHLNPHGVDRNLAWSLIYAHSDQPKTLVHHGFVSASGGHLVLDKVKKTNYPSSRSFEIGDVLTLREILDIELPALRFAG.

**Table S5. Pathogenicity assays of N-terminally GFP tagged *UhAvr1* strains showed loss of avirulence**

| Uh strains crossed | Barley cv. | No. of plants diseased (%) | No. of plants inoculated |
|---|---|---|---|
| Uh362 (*MAT-2, Uhavr1*) x Uh364 (*MAT-1, UhAvr1*) | Odessa | 32 (51) | 63 |
| | Hannchen | 0 (0) | 61 |
| Uh362 (*MAT-2, Uhavr1*) x Uh1289 (Uh364 (*MAT-1 ΔUhAvr1*)) | Odessa | 21 (46) | 46 |
| | Hannchen | 21 (43) | 49 |
| Uh362 (*MAT-2, Uhavr1*) x Uh1397 (Uh1289 (*MAT-1 ΔUhAvr1* | Odessa | 23 (43) | 53 |
| [*SP:GFP:UhAvr1*]) | Hannchen | 26 (46) | 57 |
| Uh362 (*MAT-2, Uhavr1*) x Uh1398 (Uh1289 (*MAT-1 ΔUhAvr1* | Odessa | 22 (34) | 65 |
| [*SP:GFP:UhAvr1*]) | Hannchen | 13 (19) | 69 |
| Uh362 (*MAT-2, Uhavr1*) x Uh1399 (Uh1289 (*MAT-1 ΔUhAvr1* | Odessa | 30 (54) | 56 |
| [*SP:GFP:UhAvr1*]) | Hannchen | 20 (42) | 48 |
